# Regulation of GTPase function by autophosphorylation

**DOI:** 10.1101/2021.06.23.449327

**Authors:** Christian W. Johnson, Hyuk-Soo Seo, Elizabeth M. Terrell, Fenneke KleinJan, Teklab Gebregiworgis, Genevieve M. C. Gasmi-Seabrook, Ezekiel A. Geffken, Jimit Lakhani, Kijun Song, Olesja Popow, Joao A. Paulo, Andrea Liu, Carla Mattos, Christopher B. Marshall, Mitsuhiko Ikura, Deborah K. Morrison, Sirano Dhe-Paganon, Kevin M. Haigis

## Abstract

A unifying feature of the RAS superfamily is a functionally conserved GTPase cycle that proteins use to transition between active and inactive states. Here, we demonstrate that active site autophosphorylation of some small GTPases is an intrinsic regulatory mechanism that reduces nucleotide hydrolysis and enhances nucleotide exchange, thus altering the on/off switch that forms the basis for their signaling functions. Using x-ray crystallography, nuclear magnetic resonance spectroscopy, biolayer interferometry binding assays, and molecular dynamics on autophosphorylated mutants of H-RAS and K-RAS, we show that phosphoryl transfer from GTP requires dynamic movement of the switch II domain and that autophosphorylation promotes nucleotide exchange by opening of the active site and extraction of the stabilizing Mg. Finally, we demonstrate that autophosphorylated K-RAS exhibits altered effector interactions, including a reduced affinity for RAF proteins. Thus, autophosphorylation leads to altered active site dynamics and effector interaction properties, creating a pool of GTPases that are functionally distinct from the non-phosphorylated counterpart.

## INTRODUCTION

RAS proteins and other members of the RAS superfamily coordinate cellular behaviors in response to signals. The current GTPase paradigm is that these signaling hubs are inactive when bound to guanine nucleotide diphosphate (GDP) and active when bound to guanine nucleotide triphosphate (GTP), with cycling between these states regulated by intrinsic mechanisms of GTP hydrolysis and nucleotide exchange, or by more efficient *trans* mechanisms facilitated by GTPase activating proteins (GAPs) and guanine nucleotide exchange proteins (GEFs) (Cherfils and Zeghouf, 2013). When bound to GTP, the active site of GTPases undergo conformational changes that allows interaction with, and activation of, different effector proteins. Furthermore, GTPases undergo a number of post-translational modifications (PTMs) that regulate their dynamics, subcellular localization, and activity. Dysregulation of the GTPase cycle by mutations in either GTPases or their regulators can lead to cancer, neurological diseases, or developmental syndromes (Haigis, 2017; Qu et al., 2019; Simanshu et al., 2017). While this aspect of GTPases is well studied, how disease mutations and PTM interact to modify the active state of GTPases directly is much less so.

Much of what we know about the core biochemical properties of GTPases comes from early studies of the oncogenic forms of RAS encoded by the Harvey and Kirsten RAS tumor viruses. Intriguingly, while the viral proteins exhibit a high degree of sequence identity with their cellular homologs, substitution of alanine at amino acid 59 for threonine (A59T) is the only shared difference, suggesting that the change in biochemical function resulting from this substitution provides a selective advantage for viral tumorigenesis. The Thr59 substitution is buried in the active site of RAS and undergoes autophosphorylation when RAS is bound to GTP (Shih et al., 1980). For this reason, the primary biochemical function of H-RAS was initially thought to be as a serine/threonine kinase. However, we now know that the primary biochemical function of cellular RAS proteins is GTP hydrolysis, which is defective in the viral oncoproteins due to mutations at codon 12 (G12R in v-H-RAS and G12S in v-K-RAS).

The biological advantage of Thr59 in v-RAS, and the relevance of its associated phosphorylation, is still not understood. Most members of the small GTPase superfamily have alanine at residue 59, however, a small number of family members contain a conserved autophosphorylation motif (Table 1). In some of these, as well as other GTPases where position 59 has been mutated to serine or threonine, autophosphorylation has been observed, including H-RAS (Chung et al., 1993; Chung et al., 1992; John et al., 1988), RALA (Frech et al., 1990), RAB1B (Touchot et al., 1989), and Elongation Factor Tu (Cool et al., 1990). Thus, autophosphorylation appears to be possible when either a threonine or serine nucleophile is present in the active site at this particular position, suggesting that autophosphorylation is a conserved enzymatic function in some small GTPases.

**Table 1.**
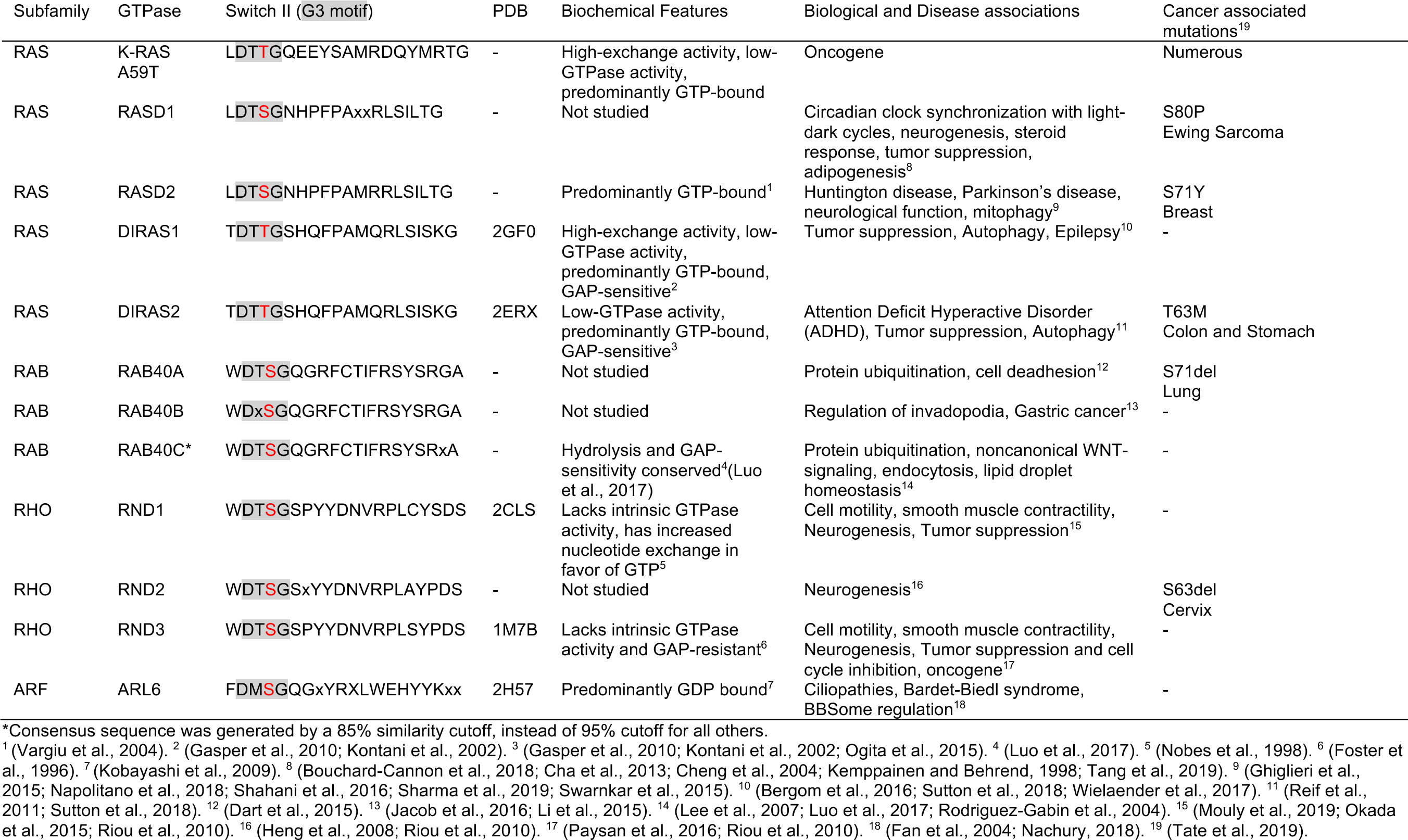
Small GTPases with autophosphorylation motif.

Mutations at Ala59 of K-, H- and N-RAS – including threonine (A59T) and glutamate (A59E) substitutions – occur in cancer, implicating autophosphorylation in RAS oncogenicity, yet the molecular and cellular properties of these mutants are poorly characterized (Haigis, 2017). In this study, we sought to understand how autophosphorylation changes small GTPase function and how this post-translational modification might influence the activity of other small GTPases. We show that the autophosphorylation mechanism is sensitive to active site dynamics, but phosphorylation itself is quite stable in solution and in cells. Using classic assays of nucleotide exchange and hydrolysis in combination with NMR, X-ray crystallography, and molecular dynamics simulations, we find that phosphorylation reorganizes the active site to favor intrinsic nucleotide exchange and inhibit GTP hydrolysis at the expense of effector interactions. Furthermore, out studies suggest that other small GTPases with an autophosphorylation motif share this unique mechanism of activation. Thus, by studying residue 59 mutants of K-RAS and H-RAS, we have discovered a new regulatory mode for the GTPase cycle that influences the functional state of GTPases with the innate ability to autophosphorylate.

## RESULTS

### Autophosphorylation alters the GTPase cycle of K-RAS

In a colon cancer cell line (SNU-175) carrying endogenous K-RAS^A59T^, we noted that K-RAS proteins showed an electrophoretic shift (Figure 1A), as did purified protein expressed in *E. coli* (Figure 1B). We confirmed by mass spectroscopy of purified protein that Thr59 was phosphorylated, indicating that phosphorylation is an intrinsic property of K-RAS^A59T^ (Figure S1A), and subsequently demonstrated in two ways that autophosphorylation resulted in the electrophoretic shift. First, purified K-RAS^A59E^ migrated at the same speed as the upper K-RAS^A59T^ band (Figure 1B). Second, the upper K-RAS^A59T^ band could be removed by lambda phosphatase treatment, albeit inefficiently and only after the protein was denatured (Figure 1C). This was an important observation, because it suggested that phosphorylated Thr59 is protected by the folded tertiary structure of K-RAS. Using the electrophoretic shift, we then measured the kinetics of autophosphorylation of purified K-RAS^A59T^ *in vitro* (Figures 1D and S1B), which produced kinetics similar to historical studies on H-RAS^A59T^, affirming that autophosphorylation occurs via an intramolecular reaction (John et al., 1988).

**Figure 1.**
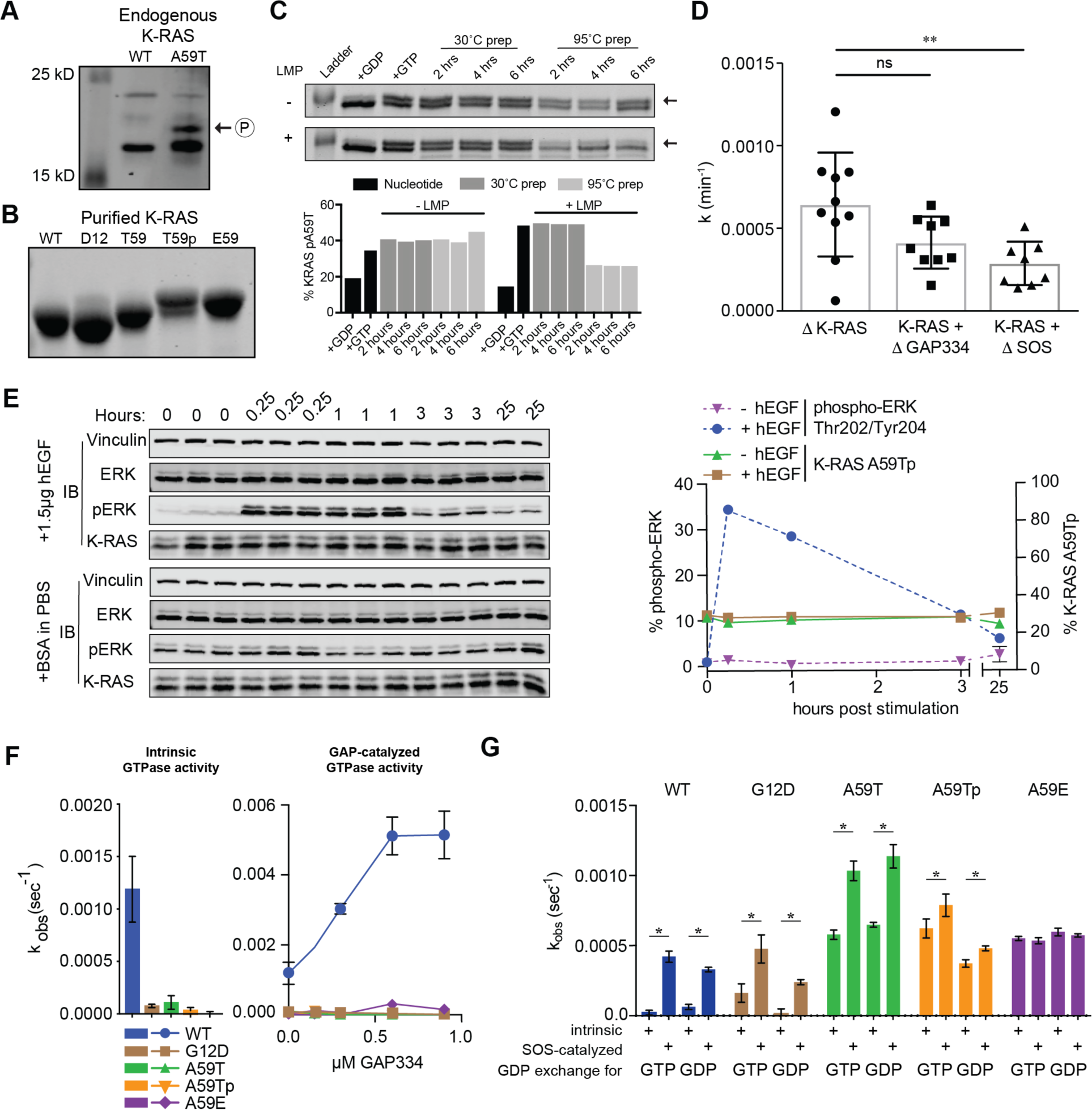
T59 phosphorylation alters K-RAS cycling. (A) Western blot of K-RAS in LIM1215 (WT) and SNU-175 (A59T) cells. (B) SOS-PAGE of purified K-RAS with different residues at position 59. (C) K-RAS^A59T^ incubated in the presence of GDP (lane 2) or GTP (lane 3). Dephosphorylation of K-RAS^A59T^ incubated with GTP with or without lambda phosphatase (LMP) (lanes 4-9) for different times at 30’C after pre-incu­ bation of protein at 30°C (lanes 4-6) or 95°C (lanes 7-9). Quantification of bands is shown below. (D) Summary of K-RAS^A59T^ autophosphorylation kinetics alone or with regulatory proteins. Effective rate constants (k) for autophosphorylation were calculated as v • [EO]^-1^ to give units of min-^1^. (E) Serum starved and hEGF (1.5µg) stimulated SNU-175 cells were analyzes for phosphorylated ERK (pERK) or K-RAS. Replicates are labelled above the gel and quantification is shown on the right. (F) Intrinsic and GAP catalyzed hydrolysis rate constants for K-RAS and mutants. Each bar or data point represents the calculated average *k* assuming a first-order mechanism (n = 3-4). (G) Measurement of exchange of GDP-loaded K-RAS4B for mant-GTP or mant-GDP. *denotes P < 0.05, and **denotes P < 0.005 by Student’s t-test.

The relationship between GTP hydrolysis and autophosphorylation has never been fully defined. One possibility is that RAS autophosphorylation is a passive byproduct of the hydrolysis reaction and Thr59 acts as an advantageous nucleophile that substitutes for the catalytic water that normally sits in the RAS active site. If this were true, we would expect GAP, which binds to RAS proteins and enhances hydrolysis, to likewise enhance the rate of autophosphorylation (Scheffzek et al., 1997). To the contrary, we found that autophosphorylation was not enhanced by GAP (Figure 1D and S1B; supplementary data). Thus, autophosphorylation is not a byproduct of hydrolysis, but is sensitive to active site conformation and is mechanistically independent of GTP hydrolysis. Furthermore, enhancement of nucleotide exchange by addition of SOS1 did not significantly affect the rate of autophosphorylation (Figure 1D and S1B; supplementary data). Since active site dynamics of K-RAS are modulated by growth signals received by cells, we next tested whether the levels of autophosphorylated K-RAS^A59T^ (K-RAS^A59Tp^) changed in response to epidermal growth factor (EGF) or insulin stimulation (Figures 1E and S1C; supplementary data). We observed no changes in the relative amount of K-RAS^A59Tp^ in these experiments, suggesting that autophosphorylation is not dynamically regulated by upstream signals.

The most compelling evidence that autophosphorylation can alter the function of a GTPase comes from the observation that A59T and A59E mutations occur in cancer. Since cancer associated mutations hyperactivate RAS proteins by altering the GTPase cycle, we examined how mutation and/or Thr59 phosphorylation affect cycling. We found that residue 59 mutants of K-RAS exhibited severely impaired intrinsic and GAP-mediated GTP hydrolysis, similar to the strongly activated mutant K-RAS^G12D^ (Figures 1F; supplementary data). The effects on nucleotide exchange were more complex (Figures 1G and S1D,E; supplementary data). First, mutation and phosphorylation strongly enhanced the rate of intrinsic exchange. Second, K-RAS^A59T^ remained sensitive to GEF-induced exchange, while K-RAS^A59Tp^ was less sensitive and K-RAS^A59E^ was entirely resistant. Third, while K-RAS^A59T^ and K-RAS^A59E^ showed no preference for GDP or GTP exchange, K-RAS^A59Tp^ preferentially exchanged GDP for GTP.

Taken together our experiments show that that K-RAS^A59Tp^ and K-RAS^A59E^ have a significant impact on the dynamics related to nucleotide exchange and hydrolysis, which likely leads to functional activation by increasing the steady state levels of K-RAS bound to GTP. We noted that many of the GTPases in Table 1 share the biochemical characteristics of residue 59 mutants, including high rates of nucleotide exchange coupled with low rates of GTP hydrolysis and dominance of the active GTP-bound state. This observation indicates that autophosphorylation is a normal feature of the functional regulation of these proteins.

### Switch II mobility promotes autophosphorylation

The mechanism of GTPase autophosphorylation is not well studied and our data suggest that it is not a byproduct of GTP hydrolysis. A previous crystal structure of H-RAS^G12V/A59T^ (PDB code 521P) shows an active site unfavorable for autophosphorylation because Thr59 is oriented away from GTP (Figure S2A) (Krengel et al., 1990). To examine the mechanism of phosphorylation, we solved two crystal structures of H-RAS^A59T^ bound to a non-hydrolyzable GTP analogue (GppNHp) (Figure S2B and Table S1). We initially chose H-RAS because it favors a closed active site, while K-RAS favors an open active site (Johnson et al., 2019; Parker et al., 2018). Active site closure is necessary for GTP hydrolysis and, while our data suggest that GTP hydrolysis and autophosphorylation occur by different mechanisms, we reasoned that active site closure was still necessary for substrate desolvation to favor phosphoryl transfer (John et al., 1993; Spoerner et al., 2010). Our crystal structures revealed that the A59T substitution disconnects switch II from both the switch I and P-loop and, in agreement with the observed reduction in GTP hydrolysis, displaces the nucleophilic water out of the active site (Figures 2A and S2C). First, Thr59 rearranges the active site by using its methyl group to push on Tyr64 and form optimal H-bonds with the backbone of Thr35 and Gln61 (black bonds (1) in Fig. 2A), resulting in switch II shifting away from the active site (black and gray bonds (2) in Fig. 2A) and breaking the H-bond between Gly60 and Gly12 in the P-loop (gray bonds (3) in Fig. 2A) that normally stabilizes active site closure upon GTP binding. Second, the Thr35-Thr59 interaction breaks the β-sheet H-bond between Thr58 and Ile36, causing destabilization of switch I. The β-sheet Thr58-Ile36 interaction is characteristic of wild-type H-RAS, but is conspicuously absent in other fast-exchange mutants of RAS, such as those with G13D mutations (Johnson et al., 2019).

**Figure 2.**
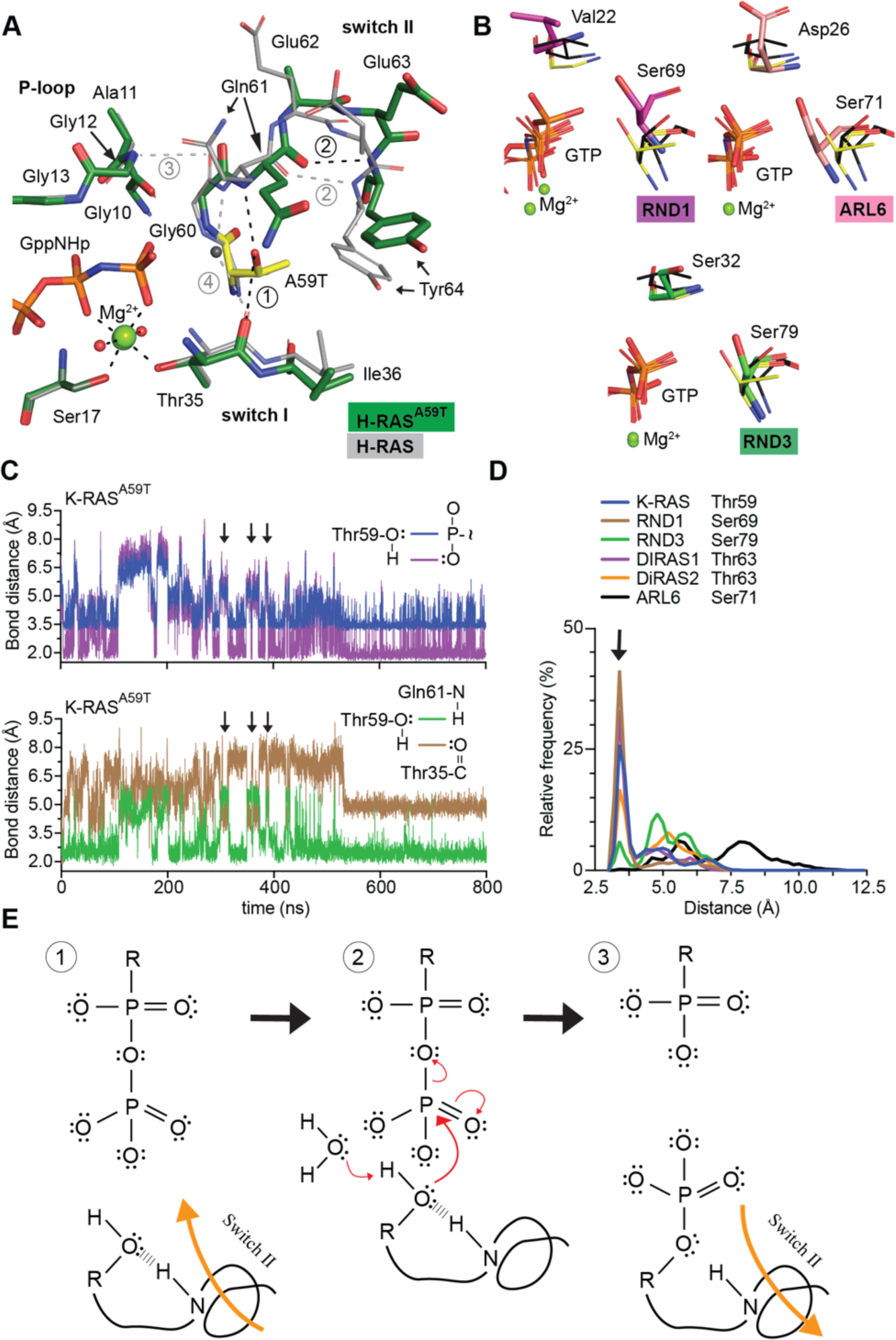
Conserved mechanism of autophosphorylation in GTPases. (A) Active site comparison of H-RAS^A59T^ crystal 1 (green) and WT H-RAS (PDB code 3K8Y,grey). Black and gray dashed lines are H-bonds made in H-RAS^A59T^ and WT structures, respectively. T59 is shown in yellow. Circled numbers are described in the text. (B) Active site similarities between H-RAS^A59T^ and other small GTPases. Black sticks are from H-RAS^G12V/A59T^ structure (PDB code 521P) with an alternate T59 orientation. (C) Bond distances made during simulation of K-RAS^A59T^ bound to GTP. Schematic insets on right show which bonds are being measured. (D) Nucleophile to gamma-phosphate distances from crystal structures of small GTPases with potential autophosphorylation activity. (E) Proposed mechanism of autophosphorylation. The N-H group of switch II represents the backbone carbonyl group of Q61.

Our H-RAS^A59T^ structures suggest that movement of Thr59 toward the γ-phosphate of GTP, a necessary step for catalysis of autophosphorylation, is possible (Figure S2D). Comparison of available crystal structures of GTP-bound small GTPases with threonine or serine at position 59 (Table 1) showed similar orientations and distances between their respective residue and the γ-phosphate, supporting their potential to undergo autophosphorylation (Figures 2B and S2E). Nevertheless, it was unclear from our data how residue 59 reaches the γ-phosphate to undergo catalysis. To answer this question, we used molecular dynamics (MD) to monitor active site motions in K-RAS^A59T^ and other potentially autophosphorylating GTPases. Throughout the K-RAS^A59T^ simulation, Thr59 closely associated with the γ-phosphate of GTP while maintaining its interaction with Gln61 (Figure 2C). In contrast, the Thr35-Thr59 interaction was commensurate with Thr59 pulling away from GTP and Q61 in switch II (Figure 2C). Consistently, most simulations of autophosphorylating GTPases showed strong association of position 59 with GTP that appeared to be regulated by residue 35 in switch I (Figures 2D and S2F).

Our analysis validates that serine/threonine nucleophiles in the proper active site position are poised for phosphoryl transfer after switch II movement toward the γ-phosphate (Figure 2E), revealing mechanistic insight into autophosphorylation of RAS and other small GTPases. The simple mechanism presented in Figure 2E is consistent with K-RAS autophosphorylation in the presence of GAP (Figure 1D). GAP makes significant interactions with switch II, essentially locking it into a particular conformation for duration of the complex and likely preventing movement of Thr59 to GTP. In contrast, the suppressive effect of SOS1 on autophosphorylation was more difficult to reconcile. First, a major determinant of the autophosphorylation reaction appears to be the distance of the nucleophile to GTP, with a decrease in distance resulting in a proportional increase in autophosphorylation (Chung et al., 1993). The structure of H-RAS bound to SOS1 shows that the exchange domain of SOS1 actually moves Ala59 toward GTP, which suggests that SOS1 should enhance autophosphorylation (Boriack-Sjodin et al., 1998), but this is not the case (Figure 1D). Instead, the decrease in K-RAS^A59T^ autophosphorylation in response to SOS1 confirms that the hydroxyl nucleophile of Thr59 must be activated and carefully oriented, likely by intramolecular interactions with switch II, to participate in catalysis.

### A ‘Mg extraction’ mechanism for hyperexchange

We next addressed how alterations at position 59 promote high rates of nucleotide exchange (hyperexchange) and why K-RAS^A59Tp^ and K-RAS^A59E^ have exchange properties that are different from K-RAS^A59T^. First, we explored the differences in dynamics between GDP bound K-RAS^A59T^ and K-RAS^A59E^ (as a mimetic of K-RAS^A59Tp^) using ^1^H-^15^N heteronuclear single quantum coherence (HSQC) NMR spectroscopy with wild-type K-RAS as a reference (Figure 3A). Consistent with our nucleotide exchange data, we observed chemical shift perturbation in backbone amide protons around the active sites of both K-RAS^A59T^ and K-RAS^A59E^ compared to wild-type K-RAS. Overall, Glu59 displayed a more global effect on K-RAS backbone chemical shifts than Thr59, indicating a more significant effect on K-RAS tertiary structure. Most notably, while the switch I region (residues 28-40) showed the largest chemical shift changes in K-RAS^A59T^, the same region experienced further peak broadening beyond detection in K-RAS^A59E^, suggesting that a conformational coupling between Thr59 and the active site in switch I is significantly enhanced by Glu59 substitution. These chemical shift changes are consistent with Thr59 motions and associated local changes around this residue in switch II, and are consistent with our H-RAS^A59T^ crystal structure and MD simulations (Figure 2).

**Figure 3.**
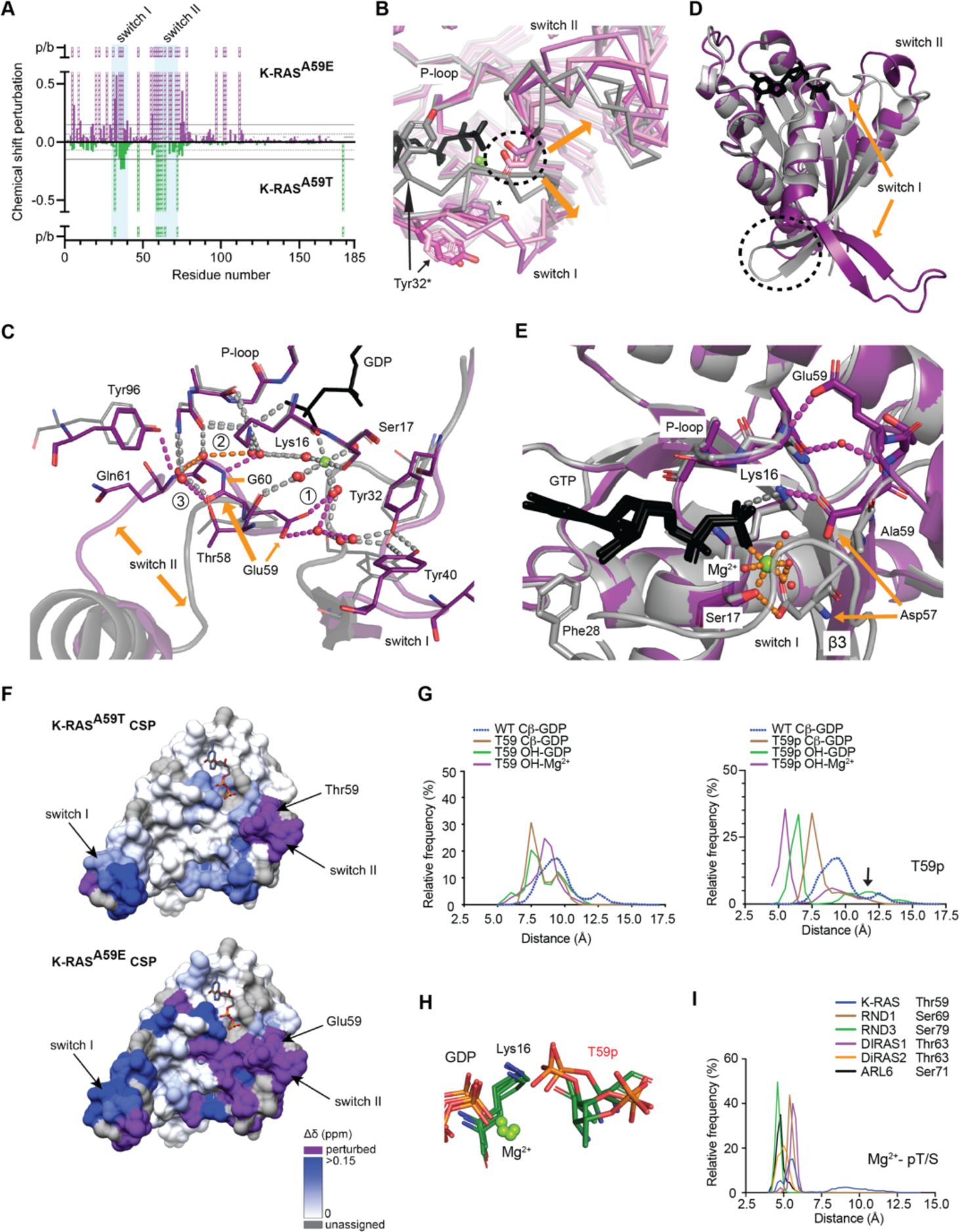
The molecular mechanism of hyperexchange. (A) Chemical shift perturbations induced by mutation of A59 plotted by K-RAS residue. Chemical shift perturbations of 1H-1SN cross-peaks for K-RAS^A59E^ or K·RAS^A59T^ are in reference to wild-type K-RAS. as descirbed in the methods section. K-RAS^A59T^ versus wild-type chemical shift perturbations are shown as negative values to compare changes to K-RAS^A59E^. Resonances t\detected in wild-type K-RAS but not in the mutants, due to large perturbations or severe peak broadening. are shown as dashed bars with an arbitrary value of 1ppm (p/b, perturbed/broadened). Residues not assigned for wild-type K-RAS are marked with a faint X. Horizontallines show threshold of mean chemical perturbation plus one standard deviation for K-RAS^A59T^ (grey. dashed) and K-RAS^A59E^ (solid black). (B) Comparison of H-RAS^A59E^ crystal structures bound to GppNHp and GDP (purple,pink) to WT H-RAS bound to GDPor GppNHp (gray). Dashed circle shows location of E59. (C) Molecule B of H-RAS^A59E^ bound to GDP (purple). E59 rearranges the active site to favor GDP release, unlike the WT reference structure (PDB code 40BE, gray).Gray dashed lines represent H-bonds shared between wildtype and K-RAS^A59E^. Colored dashes are described in the text. (D) Structure of GDP-bound K-RAS^A59E^ lacking Mg^2+^ compared to WT reference structure. Dashed circle shows junction of switch Iand helix 1. (E) E59 and D57 stabilize a pro-exchange active sitein K-RAS^A59E^. (F) Chemical shift perturbations in (A) were mapped by residue on the surface of K-RAS.The colour intensity represents increasing magnitude of chemical shift changes relative to wild-type KRAS-GDP as defined by the scale. Resonances that were detected in wild-type K-RAS but not in the mutants,due to perturbation or broadening, are shown in purple. Residues without assignments are coloured grey. The two switch regions and A59 are indicated. (G) Frequency histogram of bond distances during MDsimulations of K-RAS^A59T^ and *in silico* phosphorylated K-RAS^A59T^ bound to GDP. (H) Cluster analysis of K-RAS^A59P^ MD simulation. (I) Frequency histogram of bond distances during MD simulations of autophosphorylating GTPases bound to GDP.

To better understand the structural changes induced by autophosphorylation, and the chemical shift perturbations seen in K-RAS^A59E^, we collected crystal structures of both H-RAS^A59E^ and K-RAS^A59E^ in different nucleotide bound states (Figure S2B and Table S1). Our crystals represented different stages of nucleotide exchange induced by negative charge at residue 59. As demonstrated by H-RAS^A59E^, Glu59 repels switches I and II from the active site and weakens inter-switch β-sheet interactions, regardless of which nucleotide was bound (dashed circle in Figure 3B, Figure S3A). Moreover, Glu59 alters active site solvation to influence Mg^2+^ and GDP stability (grey dashes in Figure 3C). First, repulsion of switch I by Glu59 breaks a canonical Tyr32-Tyr40 H-bond observed in wild-type H-RAS (magenta dashes at (1) in Figure 3C), allowing Glu59 to interact with Mg^2+^, and draw switch II toward Lys16 and the P-loop (orange dashes at (2) in Figure 3C). This creates a new network that stabilizes active site opening and draws Lys16 and Mg^2+^ away from GDP (magenta dashes at (3) in Figure 3C).

The impact of Mg^2+^ release is demonstrated by the active site of K-RAS^A59E^, which appeared to be an intermediate of intrinsic exchange and which was similar to the crystal structure of K-RAS^A146T^, a mutant that exhibits fast GEF-independent nucleotide exchange (Poulin et al., 2019). The active site of K-RAS^A59E^ is completely open and lacks Mg^2+^, with switch I and β2 peeled away from the globular domain to form a novel β-sheet (orange arrow in Figure 3D). Mg^2+^ dissociation allows β3 to extend toward switch II and Glu59 and Asp57 to make salt-bridges with the P-loop and K16 respectively (magenta dashes in Figure 3E). From the novel beta sheet, β2 leads into the interswitch loop 3 with rearrangement of salt-bridges between Arg164, Asp47, and Glu49 at the junction of switch I and helix 1 (dashed circle in Figures 3D and S3B). Ultimately, these changes remove Pi-stacking interactions between Phe28 in switch I with the guanine base of GDP, increasing solvent exposure of the nucleotide (Figures 3E and S3C). The details of our H-RAS^A59E^ and K-RAS^A59E^ crystal structures are consistent with peak broadening in switches I and II seen of our HSQC NMR data, and this was particularly obvious after mapping our chemical shift changes onto the crystal structure of K-RAS^A59E^ (Figure 3F).

Our data suggest a mechanism of ‘Mg^2+^ extraction’ that supports hyperexchange by autophosphorylated K-RAS^A59T^ and, possibly, other autophosphorylating GTPases. To explore this possibility, we used MD simulations to examine the dynamic relationship of position 59, Mg^2+^, and GDP in RAS proteins and other GTPases. Residue 59 Cβ atom motions in K-RAS^A59T^ showed that both mutation and the charge associated with phosphorylation shift residue 59 and switch II into the active site (Figures 3G,H and S3D). These results were consistent with simulations for other autophosphorylating GTPases, which also showed that phosphorylated position 59 moves toward the respective GDP and Mg^2+^ (Figures 3I and S3E). Together, our MD simulations and crystal structures show that negative charge at position 59 destabilizes the Mg^2+^ - GDP interaction.

For RAS in particular, ‘Mg^2+^ extraction’ appeared to represent an intermediate step in the process of taking on the conformation seen in our crystal structure of K-RAS^A59E^, as we noted that the overall changes in K-RAS structure and dynamics seen in our NMR experiments were more consistent with the K-RAS^A59E^ crystal structure than the H-RAS^A59E^ crystal structure (Figure 3F). This last point also explains why K-RAS^A59Tp^ and K-RAS^A59E^ have different overall kinetics of nucleotide exchange compared to K-RAS^A59T^. While the rate of intrinsic nucleotide exchange of GDP for GTP is essentially the same for K-RAS^A59T^, K-RAS^A59Tp^, and K-RAS^A59E^, only the non-phosphorylated protein has a considerable response to SOS1. Thus, while K-RAS^A59T^ locally shifts switch I and II away from the nucleotide to enhance exchange (upper vs lower panel, Figure 3F), the overall conformation is likely still recognizable and sensitive to SOS1 interaction. In contrast, the extended and open active site conformation of K-RAS^A59E^, and presumably K-RAS^A59Tp^, is resistant to recognition and complex formation by SOS1. Although our NMR experiments do not provide direct evidence for the β-sheet structure seen in the K-RAS^A59E^ crystal, that Glu59 stabilizes K-RAS with an extended and open active site conformation is consistent with two other crystal structures, K-RAS^A146T^ and wild-type K-RAS, that have this same structural feature (Dharmaiah et al., 2019; Poulin et al., 2019). Dharmaiah et al. suggested that the active site features of these K-RAS crystals are the result of either a missing initiator methionine or N-terminal acetylation. However, they were unable to show that methionine and N-acetylation alters the activity of SOS1 with wild-type K-RAS or chemical shift changes in switch I by HSQC NMR. Likewise, the nucleotide exchange rate of K-RAS^A146T^ is enhanced by SOS1, unlike K-RAS^A59Tp^ and K-RAS^A59E^. Taken together, these observations suggest that the extended and open conformation is likely present in wild-type K-RAS, but is less favored and in conformational equilibrium with other active site states. In contrast, Glu59 favors and stabilizes the extended and open conformation in K-RAS^A59E^ through additional salt-bridges (Figure 3E). This interpretation also suggests that the active site of RAS only needs to open so much for intrinsic exchange of GDP for GTP, as K-RAS^A59T^, K-RAS^A59Tp^, and K-RAS^A59E^ all converge on similar rate constants for this reaction (Figure 1G), and that the mechanism of SOS1-catalyzed nucleotide exchange is similar but not identical to the intrinsic mechanism of nucleotide exchange. Thus, our different biophysical and theoretical approaches argue that introduction of a Ser or Thr functional group at ‘residue 59’ alters the enzymatic function of small GTPases to enable autophosphorylation, which permanently alters active site dynamics to favor intrinsic nucleotide exchange.

### Autophosphorylation activates K-RAS

The active site dynamics of K-RAS^A59Tp^ and K-RAS^A59E^ that are necessary for nucleotide exchange appear to be at odds with GTPase function, as small GTPases require closure of the active site to interact with their known effectors (Lu et al., 2016; Vetter, 2017). Nevertheless, the fact that codon 59 mutations occur in cancer suggest that they functionally activate K-RAS. To address this paradox, we first examined K-RAS^A59T^ function in SNU-175 cells expressing a doxycycline inducible short-hairpin RNA (shRNA) targeting the *KRAS* transcript. Knockdown of K-RAS in SNU-175 cells reduced proliferation and ERK phosphorylation, indicating that the A59T mutation does promote a proliferative function of K-RAS (Figure 4A-B and S4A). At the same time, knockdown of K-RAS caused a relative increase in K-RAS^A59Tp^ (Figure 4C). This result confirms the independent and passive nature of K-RAS^A59T^ autophosphorylation, as the phosphorylated form of v-H-RAS persists in the cell for a longer period of time than the non-phosphorylated form (Ulsh and Shih, 1984).

**Figure 4.**
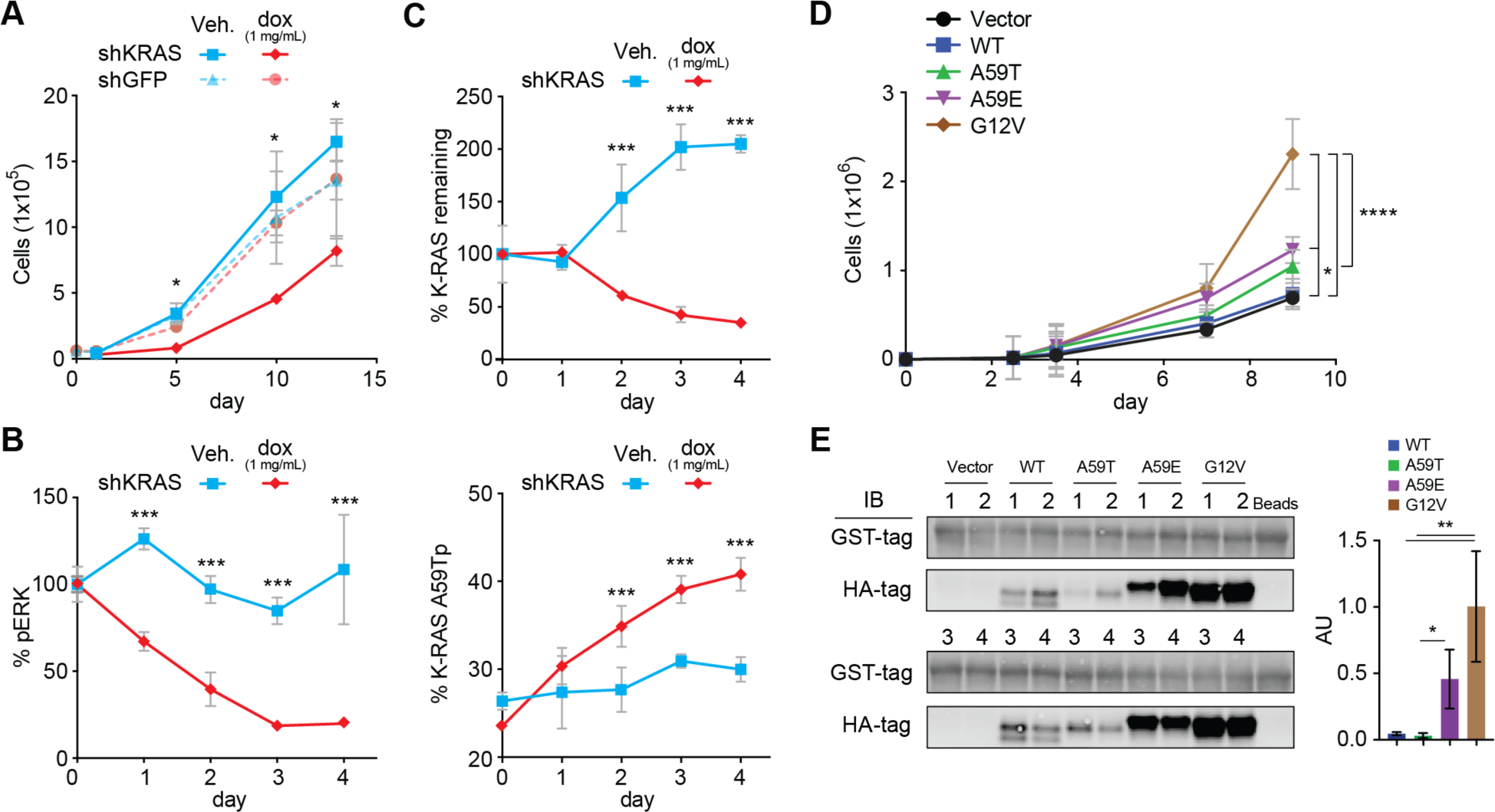
Role of KRAS and KRAS in cell transformation. (A) Effect of *KRAS* shRNA on proliferation in SNU-175 cells (n=2-3). (B) Effect of *KRAS* shRNA on ERK phosphorylation in SNU-175 cells (n=2-3). (C) Effect of shRNA on K-RAS expression and autophosphoryation in SNU-175 cells (n=2-3). (C) Proliferation of NIH3T3 cells in 6% serum ectopically expressing HA-K-RAS (n=4). (D) Precipitation of GTP bound HA-K-RAS mutants by C-RAF-RBD-GST coupled to glutathione-agarose. Quantitation on the right.Statistical analysis was performed with students T-test. *, **, ***, **** represent P-values of <0.05, <0.005, and <0.0005.

While knockdown of K-RAS in the SNU-175 cells demonstrated dependence of this line on mutant K-RAS, we were unable to evaluate the relative activity of different K-RAS mutants. We next tested the impact of ectopic expression of Ala59 mutants in NIH3T3 fibroblasts (Figure S4B). Ectopic expression of K-RAS^A59T^ and K-RAS^A59E^ weakly enhanced cell proliferation compared to K-RAS^G12V^, a common oncogenic mutant (Figure 4D). This observation was surprising given the strong effect that A59T/E have on GTP hydrolysis and nucleotide exchange (Figure 1F,G). To understand these differences in fibroblast transformation, we measured the ability of these proteins to interact with the RAS binding domain of C-RAF (C-RAF-RBD), which recognizes and binds to GTP-bound RAS and provides a readout for the active state of these proteins. In contrast to our expectation that K-RAS^A59T^ and K-RAS^A59E^ would be similarly activated compared to K-RAS^G12V^, both the K-RAS^A59T^ and K-RAS^A59Tp^ proteins failed to interact with C-RAF-RBD, while the K-RAS^A59E^ was generally attenuated (Figure 4E). Together, these data demonstrate that K-RAS^A59T/E^ can transform cells, but that their oncogenic activity is potentially limited by active site dynamics that might negatively impact effector interactions.

### Autophosphorylation influences K-RAS effector interactions

Our data indicated that residue 59 phosphorylation might inhibit effector binding due to its influence on switches I and II, which constitute effector binding interfaces (yellow surfaces in Figure 5A). To test this idea, we preloaded mutant K-RAS protein in our NIH3T3 lysates with GTP and again tested them for interaction with C-RAF-RBD or, additionally, with full-length RASSF5 protein. These effector interactions provide complementary information because C-RAF-RBD interacts exclusively with switch I, while RASSF5 interacts with both switch I and II. We found that T59 phosphorylation inhibits interactions of C-RAF and RASSF5, while A59E selectively inhibits interaction with RASSF5 (Figure 5B).

**Figure 5.**
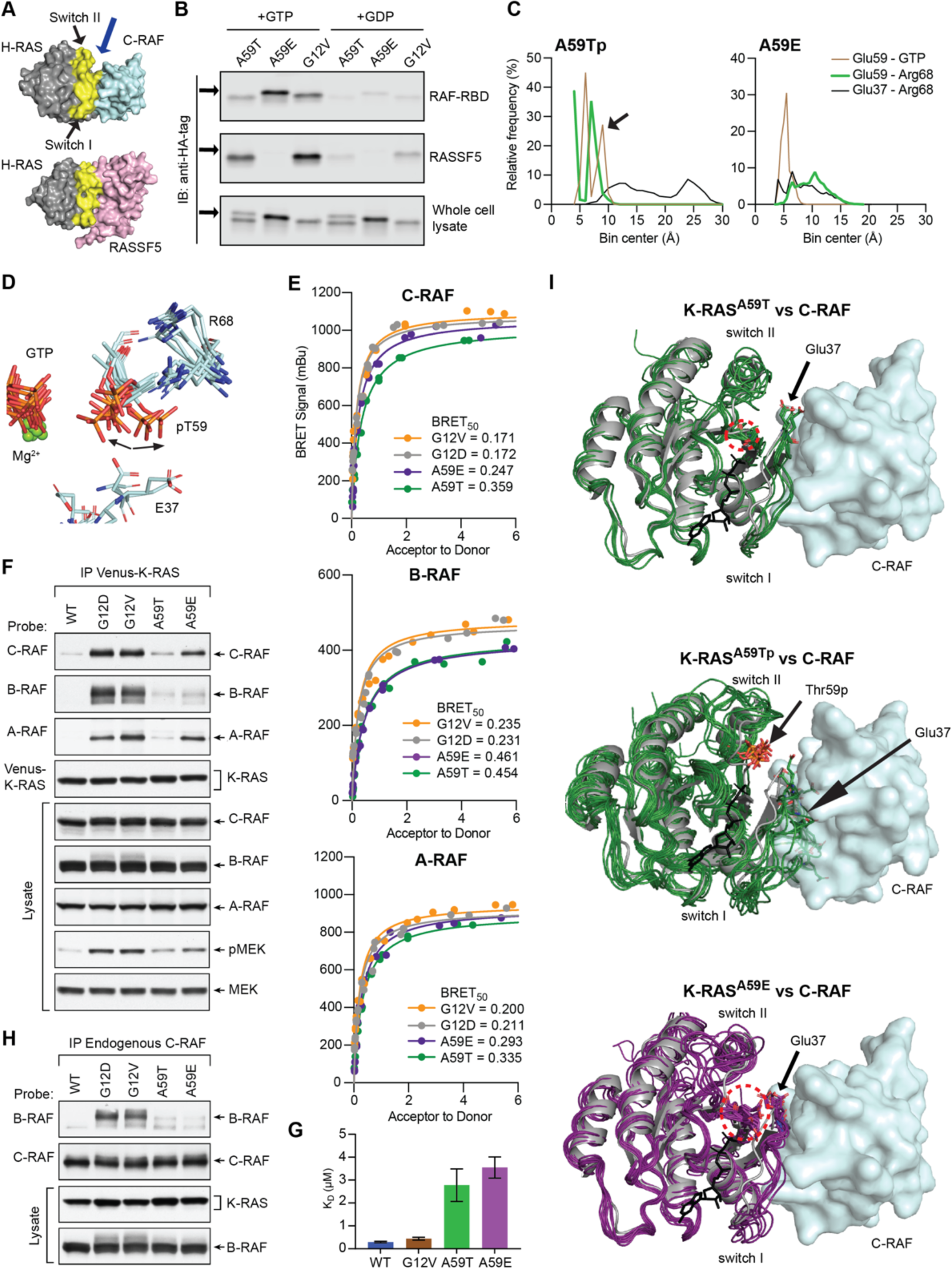
GTP induces phosphoryl movement to discriminate effector interactions. (A) Effector binding interfaces of C-RAF-RBD (PDB code 4G0N) and RASSF5-RA (PDB code 3DDC) domains. Blue arrow shows location of A59. (B) Pulldown of ectopic HA-KRAS4B, preloaded with GTP or GDP,by GST-C-RAF-RBD or GST-RASSF5. (C) Bond distances during simulation of mutant K-RAS bound to GTP. (D) Simulation cluster analysis of phosphorylated K-RAS^A59T^. (E) BRET saturation curves showing the interaction of RAF isoforms with the indicated K-RAS mutants in HEK293FT cells. (F) Western blot of RAF isoforms coimmunoprecipitating with Venus-tagged K-RAS proteins from serum-starved HeLa cells. (G) Affinities of GppNHp bound K-RAS (wild-type, G12V, A59E and A59T mutants) for the RBD of B-RAF determined by BU. Error bars represent standard deviation (n = 2-4). (H) Co-immunoprecipitation of B-RAF and C-RAF with different K-RAS mutants from serum-starved Hela cells. Also shown is dimerization of B-RAF and C-RAFinduced by the K-RAS mutants. (I) Comparison of K-RAS proteins (green) conformations generated by cluster analysis of MD simulations to H-RAS (gray) bound to C-RAF (cyan). Nucleotide is shown as black sticks.

Since our GppNHp bound H-RAS^A59E^ crystal structures have disordered active sites, we used MD simulations to explain our binding results, in particular why K-RAS^A59Tp^ and K-RAS^A59E^ appeared to behave differently. We had noted that simulations of GDP-bound K-RAS^A59Tp^ exhibited a low frequency conformation of the phosphoryl group facing away from the active site (arrow in Figure 3G,H). Likewise, while wild-type K-RAS and K-RAS^A59T^ had similar overall dynamic profiles (Figure S5A,B), the phosphoryl group of K-RAS^A59Tp^ showed a clear transition out of the active site, causing repulsion of Glu37 (arrows in Figure 5C,D). In contrast, K-RAS^A59E^ did not make this transition (Figures 5C and S5C), and, in fact, appeared to enhance switch I-II interactions to some degree, as Glu37 in switch I and R68 in switch I, directly interact during the simulation. With the exception of ARL6 and DIRAS1, MD simulations suggested that phosphoryl-transitions in the other autophosphorylating GTPases, and their overall effects on switches I and II in the models, were similar to phosphorylated K-RAS^A59T^ (Figure S5D).

In light of these findings, the GTP-bound state of K-RAS^A59T^ and K-RAS^A59E^ was likely underestimated in our initial RAS activity assays because autophosphorylation impacts the affinity of RAS-RAF interaction. To address this directly, we tested RAF binding activity of K-RAS^A59T^ and K-RAS^A59E^ using bioluminescence energy transfer (BRET) in cells (Terrell et al., 2019). As in our binding experiments with C-RAF-RBD, K-RAS^A59T^ and K-RAS^A59E^ exhibited reduced affinity for full length C-RAF, as well as A-RAF and B-RAF, compared to the common oncogenic mutants K-RAS^G12V^ and K-RAS^G12D^ (Figure 5E,F and supplementary data). The reduced interaction with B-RAF is likely due to a significantly reduced affinity for B-RAF in solution, due to both a reduction in kon and increase in koff for K-RAS/B-RAF complex formation (Figures 5G and Figure S6A-D,H). Furthermore, the status of MEK phosphorylation in cells expressing K-RAS^A59T^ or K-RAS^A59E^ trended with C-RAF co-immunoprecipitation (Figure 5F), reflecting the graded differences in fibroblast proliferation induced by K-RAS^A59T^ and K-RAS^A59E^ (Figure 4C). While B-RAF has a clear change in association with K-RAS^A59T^ and K-RAS^A59E^, it could still be activated through a K-RAS/C-RAF interaction, which could subsequently induce heterodimerization with B-RAF. We tested this possibility, but found that neither K-RAS^A59T^ and K-RAS^A59E^ promoted C-RAF/B-RAF heterodimerization to a similar extent as K-RAS^G12D^ or K-RAS^G12V^ (Figure 5H). Together, our data suggest that K-RAS^A59T^ and K-RAS^A59E^ do not activate the MAPK signaling through B-RAF, but rather through weak activation of A-RAF and C-RAF.

The reduction in RASSF5 binding in response to codon 59 mutation was also noteworthy. RASSF5 is a member of putative tumor suppressor family (RASSF1-10) that is associated with growth inhibition downstream of activated RAS (Volodko et al., 2014). We discovered that only RASSF1 and RASSF5 have significant affinity for active K-RAS and that K-RAS^A59Tp^ and K-RAS^A59E^ appeared to specifically affect the interaction with RASSF5 (Figure S6E-G,H and Figure S7A-B). The functional activation of K-RAS – *i.e.* its induction of proliferation – by codon 59 mutations might be due to its weak activation of pro-proliferation signaling and its lack of activation of anti-proliferation signaling.

For small GTPases, increased active site dynamics in the GTP-bound state, along with phosphorylation that changes active site compaction and organization, will necessarily alter effector affinity and kinetics of complex formation. This is demonstrated by comparison of our MD simulations to crystal structures of H-RAS bound to C-RAF-RBD (Figure 5I) and RASSF5 (Figure S7C). For instance, K-RAS^A59T^ shows weak affinity for both B-RAF and RASSF5 (Figure 5G and Figure S7A). For B-RAF, this makes sense because our MD simulations of K-RAS^A59T^ show that Thr59 association with GTP is disruptive to Thr35 packing in the active site (Figure 2C and Figure S2F), and it is well known that Thr35 mutants of K-RAS and H-RAS have reduced affinity for C-RAF (Hamad et al., 2002). For the K-RAS^A59T^/RASSF5 interaction, weakened affinity can be explained by the expanded binding interface of RASSF5, which is likely more sensitive to active site compaction necessary for a strong complex to form (Stieglitz et al., 2008). After Thr59 phosphorylation, dynamics in switch I and II increase significantly, preventing either C-RAF, presumably B-RAF, or RASSF5 binding to K-RAS^A59Tp^ (Figure 5I and Figure S7C). Increase in switch II dynamics is likely also a significant reason why K-RAS^A59E^ cannot bind RASSF5. However, K-RAS^A59E^ was more competent than K-RAS^A59T^ in binding C-RAF (Figure 5E,F), likely due to the Glu59 having less dynamical changes in switch I, while changes in switch II are less consequential. Movement of Glu37 in our MD simulations of K-RAS was also notable and helps explain our data, because Glu37 mutants, like Thr35, have reduced affinity for C-RAF (Hamad et al., 2002). Furthermore, mutation of Glu37 to glycine improves binding of RAS for RASSF5 (Khokhlatchev et al., 2002). Thus, repulsion of Glu37 by Glu59, or phosphorylated Thr59, further inhibits binding of RASSF5 to K-RAS^A59E^ and K-RAS^A59Tp^ (Figure 5B, and Figure S7A,B). These data demonstrate a significant consequence of GTPase autophosphorylation is to alter effector engagement.

## Discussion

Here, we elucidate the mechanistic basis for the autophosphorylation of small GTPases and determine the effect of this modification on the GTPase cycle. The functional state of a GTPase is presumed to be a function of its nucleotide binding state (Figure 6). In the canonical GTPase cycle, intrinsic and GEF-induced nucleotide exchange activate the protein by loading GTP into the active site, while intrinsic and GAP-induced GTP hydrolysis serve to inactivate. Using K-RAS^A59T^ as an architype for small GTPases that are capable of autophosphorylation, we show that phosphorylation at position 59 alters active site dynamics, nucleotide exchange, and effector interaction to dramatically reorganize the GTPase cycle (Figure 6). Although the ability of H-RAS^A59T^ to autophosphorylate has been known for some time, our study disproves one of the initial presumptions of autophosphorylation – that it is simply a byproduct of the hydrolysis reaction (John et al., 1988). We found that binding of GAP to K-RAS^A59T^ actually slows autophosphorylation (Figure 1D), confirming that autophosphorylation occurs via a different catalytic mechanism.

**Figure 6.**
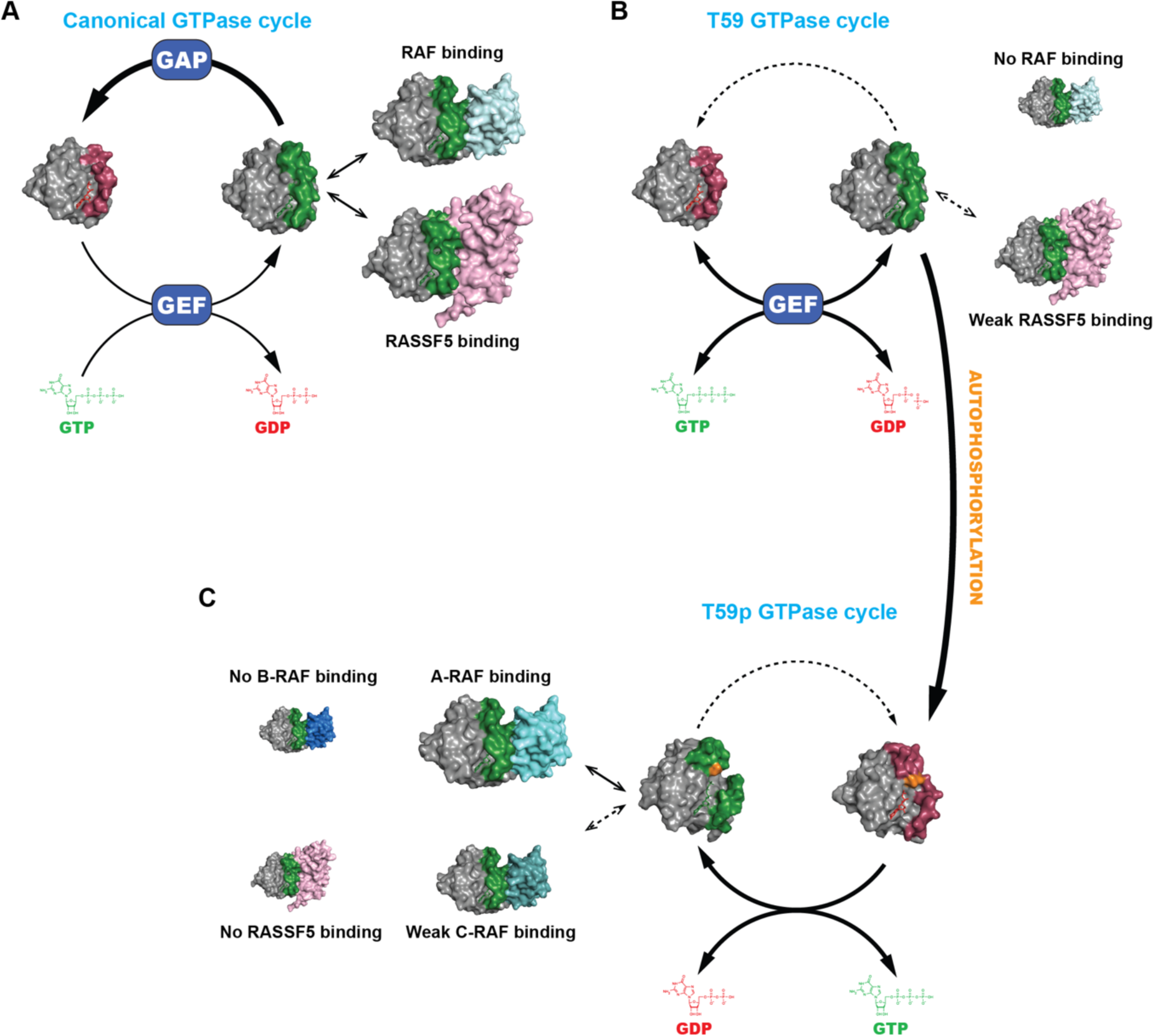
Influence of autophosphorylation on nucleotide cycling and effector engagement. (A) In a normal cell, K-RAS activity is low due to increased activity of GAPs relative to GEFs, pushing the equilibrium toward the GDP-bound state. When K-RAS is bound to GTP, it can engage a variety of different effectors. (B) Mutation of alanine 59 to threonine inhibits nucleotide hydrolysis and promotes intrinsic nucleotide exchange, creating an activated form of K-RAS. When bound to GTP, the mutant does not interact with RAF proteins and exhibits weak binding to RASSF5. (C) When threonine 59 becomes phosphorylated (orange), K-RAS enters an alternative cycle where it loses sensitivity to GEF. When bound to GTP, phosphorylated K-RAS retains some RAF binding, but loses the ability to bind B-RAF and RASSF5.

An intriguing outcome of our study is that autophosphorylation enhances intrinsic exchange but desensitizes K-RAS to SOS1-catalyzed nucleotide exchange (Figure 1G). Moreover, phosphorylation of Thr59 creates a preference for GTP exchange over GDP exchange (Figure 1G). After GTP binding, K-RAS^A59Tp^ and K-RAS^A59E^ do undergo necessary dynamics to take on an active GTP-bound state, but with altered effector binding (Figure 5). Altered interaction with RAF proteins likely limits pro-proliferation signaling downstream of K-RAS^A59T^ and K-RAS^A59E^. It is also possible that autophosphorylation allows for novel effector interactions, perhaps overlapping with the other small GTPases that potentially autophosphorylate (Table 1). Indeed, DiRas1/2 are RAS subfamily members can bind and compete with K-RAS effectors (Bergom et al., 2016). One question that arose from our early observations is why codon 59 mutations are so rare in cancer when they impact GTP hydrolysis to the same extent as very common codon 12 mutants. Codon 59 mutations also increase nucleotide exchange and, although increasing nucleotide exchange can typically activate RAS proteins (Haigis, 2017), functional activation of K-RAS^A59T/E^ is likely limited by the mechanism of hyperexchange and its altered effector interactions, ultimately creating weakly activated mutants. This attenuating function of Thr59 might be necessary for v-K-RAS and v-H-RAS to avoid oncogene-induced senescence due to hyperactivated MAPK signaling (Li et al., 2018).

K-RAS^A59T^ is a somatic mutant that occurs in cancer, however, several GTPases share the ability to autophosphorylate when mutated to serine or threonine at the correct active site position (Chung et al., 1993; Chung et al., 1992; Cool et al., 1990; Frech et al., 1990; John et al., 1988; Touchot et al., 1989). Since the field has yet to identify GEFs for many of the GTPases listed in Table 1, it is possible that threonine or serine at position 59, as well as autophosphorylation, might instead provide this regulatory role. For GTPases capable of autophosphorylation, this altered nucleotide binding cycle is likely an aspect of their normal regulation, creating a pool that is functionally distinct from the non-phosphorylated pool. We also noted that some of these small GTPases have cancer associated mutations at their analogous residue 59 (Table 1), which would render them unable to auto-phosphorylate, and perhaps altering their GTPase cycle back toward the more canonical form.

The most thoroughly characterized RAS modifications (*e.g.* phosphorylation, di-ubiquitination) regulate its subcellular localization, typically pushing it from the plasma membrane to organellar endomembranes (Ahearn et al., 2018). But others (*e.g.* mono-ubiquitination, acetylation, AMPylation, tyrosyl phosphorylation) are reported to affect the GTPase cycle by influencing nucleotide exchange (Barthelmes et al., 2020; Kano et al., 2019; Sasaki et al., 2011; Yang et al., 2012). Indeed, our work has parallels with a recent study showing that Src shifts K-RAS into a ‘dark state’ through phosphorylation of Tyr32 and Tyr64. These modifications stabilize the GTP-bound state of K-RAS while simultaneously preventing K-RAS from interacting with RAF kinases (Kano et al., 2019). However, this study differs from that of Kano et al. in that phosphorylation of residue 59 still promotes functional activation of K-RAS. Thus, our work illuminates a new paradigm for regulation of the GTPase cycle via autophosphorylation. This discovery opens the door to a deeper look at the PTM landscape on GTPases and more detailed studies of their functional ramifications.

## Methods

### Statistics statement

No statistical methods were used to predetermine sample size. The experiments were not randomized, nor were investigators blinded to sample allocation and experimental assessment. For biochemical or enzymology experiments, t-tests were used to determine significance. For biological experiments, the Mann-Whitney U-test was used to determine significance.

Sequence alignment of small GTPases with autophosphorylation motif *S*equences for K-RAS, ARL6, DIRAS1, DIRAS2, RAB40A, RAB40B, RAB40C, RASD1, RASD2, RND1, RND2, and RND3 were extracted from the NCBI database (Coordinators, 2016). Alignments were done using the T-COFFEE algorithm provided by SnapGene® software (GSL Biotech).

### Mass spectrometry

Approximately 20 micrograms of purified 6xHis-tagged KRAS4B^A59T^ were reduced with 5 mM tris(2-carboxyethyl)phosphine (ThermoFisher Scientific, Cat #77720) for 30 min at room temperature, alkylated with 10 mM iodoacetamide (ThermoFisher Scientific, Cat #A39271) for 30 min at room temperature in the dark, and quenched with 10 mM dithiotreitol for 15 min at room temperature. The sample was divided into two replicates, precipitated by trichloroacetic acid precipitation and digested with 1:50 (w/w) chymotrypsin (Promega, Cat #V106A) in 100 mM Tris-HCl and 10 mM CaCl2 for 18 h at 37 °C with shaking. The samples were desalted using a StageTip, dried by vacuum centrifugation and solubilized in 5% acetonitrile and 5% formic acid. LC-MS/MS analysis was performed on a Q Exactive mass spectrometer (Thermo Fisher Scientific) coupled with a Famos Autosampler (LC Packings) and an Accela600 liquid chromatography pump (Thermo Fisher Scientific). Peptides were separated on a 100 μm inner diameter microcapillary column packed with ∼25 cm of Accucore C18 resin (2.6 μm, 150 Å, Thermo Fisher Scientific). For each analysis we loaded ∼1μg onto the column.

Peptides were separated using a one-hour method of 5 to 25% acetonitrile in 0.125% formic acid with a flow rate of ∼300 nL/min. The scan sequence began with an Orbitrap MS^1^ spectrum with the following parameters: resolution 70,000, scan range 300−1500 Th, automatic gain control target 1 × 10^5^, maximum injection time 250 ms, and centroid spectrum data type. We selected the top twenty precursors for MS^2^ analysis which consisted of high-energy collision dissociation with the following parameters: resolution 17,500, AGC 1 × 10^5^, maximum injection time 60 ms, isolation window 2 Th, normalized collision energy 30, and centroid spectrum data type. The underfill ratio was set at 1%, which corresponds to a 1.1 × 10^4^ intensity threshold. Unassigned and singly charged species were excluded from MS^2^ analysis and dynamic exclusion was set to automatic.

Spectra were searched using Sequest with a 3 ppm precursor mass tolerance, 0.03 fragment ion tolerance, and no limit on internal cleavage sites (Eng et al., 1994). Cysteine alkylation was set as a fixed modification, methionine oxidation and phosphorylation of serine, threonine, and tyrosine were included as variable modifications. Spectra were searched against a custom database containing the 6xHis-KRAS4B^A59T^ sequence, common contaminants, and the reversed peptide sequences. False discovery rate was estimated by linear discriminant analysis and applied at one percent at the peptide level (Elias and Gygi, 2007; Peng et al., 2003).

### Molecular dynamics simulations

Starting coordinates for molecular dynamic simulation were generated from the crystal structures published here and from the PDB. For simulations of GDP-bound or GTP-bound K-RAS, we used the starting wild-type structures 6MBU (Dharmaiah et al., 2019) and 4DSO (Maurer et al., 2012). Starting PDB coordinates for RND1, RND3, DIRAS1, DIRAS2, and ARL6 were 2CLS, 1M7B (Fiegen et al., 2002), 2GF0, 2ERX, and 2H57, respectively. Starting structures went through an extra round of refinement or modeling to add in missing residues and remove alternate conformations. Structures were converted into the GDP- or GTP-bound states, or mutated, *in silico*. Preparation of starting files, including ‘residue 59’ phosphorylation was done using ‘solution builder’ available from CHARM-GUI (Brooks et al., 2009; Jo et al., 2008; Lee et al., 2016). Charged residues, including protein termini, were protonated or deprotonated in accordance with neutral pH. A cubic box with edges 10Å from each protein was created and filled with TIP3P water molecules and neutralized with Cl^-^ and Na^+^ ions to 150mM. Minimizations, equilibrations, and simulations were done using GROMACS (ver. 2020.1) and a GPU server featuring 8x Tesla v100 workstation, on the O2 High Performance Compute Cluster, supported by the Research Computing Group, at Harvard Medical School. Solvated systems were energy minimized by steep integration for 5000 steps or at a maximum force of 1000 kJ/mol/nm or less. The Verlet cutoff scheme was used for nonbonded atoms and the LINCS algorithm was applied to covalent H-bonds. Short-range van der Waals interactions were switched off from 1.0-1.2 nm, and long-range interactions were computed using the Particle Mesh Ewald method. Simulation temperatures were maintained at 310K using Nose-Hoover extended ensemble. The isothermal-isobaric ensemble (NPT) was generated using the Parrinello-Rahman barostat method with periodic boundary conditions. Simulations were done for 800-900ns using the GROMOS force-field. Validation and analyses steps were done in GROMACS, including cluster analysis using the GROMOS algorithm (Daura et al., 1999). Distance measurements and visual analyses were done using PyMOL and VMD (Humphrey et al., 1996).

#### Protein purification for biochemical assays

Human K-RAS4B and the catalytic domains of GAP (715-1047) and SOS11 (564-1049) were synthesized with N-terminal hexa-histidine tags (6xHis) and placed in the pET21a plasmid for *E. coli* expression by GENEWIZ. Expressed K-RAS proteins were full-length with the C-terminal two amino acids removed to mimic post-translational processing (Hancock et al., 1991). Protein expression was done using chemically transformed BL21 (DE3) strain of *E. coli* and Terrific Broth media. A 30mL culture of broth was inoculated with BL21 E. coli cells with the appropriate tagged protein and allowed to grow overnight at 37°C with agitation. The following morning 25mL of culture was applied to a 1L culture of terrific broth, and protein expression was induced with 120mg/L of IPTG after *E. coli* reached an O.D. 0.8. After 6 hours at 37°C the cell solution was centrifuged at 10,000RPM at 4°C and pellets were stored at -80°C. His-tagged protein was purified from *E. coli* paste as follows. Frozen pellets were re-suspended in sterile filtered buffer TA (20mM Tris pH 7.5, 50mM NaCl, 5mM MgCl2, 5% glycerol, 10µM GDP, 156µL/L 2-mercaptoethanol) containing 4mM PMSF at <0.5mg/mL by agitation at room temperature for 15-30 minutes. E. coli cells were lysed by sonication on ice. Lysate was then clarified for 30 minutes at 4°C and 14,000 RPM. Chromatography of supernatant was done using an AKTA FPLC (GE healthcare sciences). Supernatant was first subjected to immobilized metal affinity chromatography (IMAC) using a 5mL Hi-Trap TALON column (GE healthcare sciences). The running buffer for IMAC was buffer TA, and sterile buffer TB (20mM Tris pH 7.5, 50mM NaCl, 5mM MgCl2, 200mM Imidazole, 5% glycerol, 10µM GDP, 156µL/L of 2-mercaptoethanol) was used for gradient elution. Post-elution, positive fractions were screened by SDS-PAGE, pooled and concentrated to <1mL using a 10,000 kDa Amicon Ultra spin concentrator (Fisher Scientific). Concentrated protein was then buffer exchanged into QA buffer (20mM Tris pH 8.0, 50mM NaCl, 5mM MgCl2, 5% glycerol, 10µM GDP, 156µL/L 2-mercaptoethanol) using a ZEBA spin column and its recommended protocol (Thermo Fisher Scientific). Buffer exchanged protein was applied to a 5mL HiTrap QHP column (GE Healthcare,) using a 30% gradient of QB buffer (20mM Tris pH 8.0, 1M NaCl, 5mM MgCl2, 5% glycerol, 10µM GDP, 156µL/L 2-mercaptoethanol) over 60mL. Positive fractions were then pooled and subjected to buffer exchange twice using ZEBA spin columns. The first exchange was into protein into stabilization buffer (20mM Tris pH 7.5, 10mM NaCl, 1mM MgCl2, 1% glycerol, 1mM DTT) containing 5mM EDTA and then protein was re-exchanged into stabilization buffer without EDTA. Protein concentration was then determined, and equimolar GDP was added to KRAS4B protein. Protein for nucleotide hydrolysis and exchange were diluted to approximately 100µM, including GAP334 and SOS1cat, flash frozen in 50-150µL aliquots for single use, and then stored at -80°C. To obtain phosphorylated K-RAS4B, after the IMAC step, K-RAS4B^A59T^ was incubated overnight or longer in the presence of GTP, and then purification was continued. Post-QHP, phosphorylated protein was identified by SDS-PAGE and pooled separately from the non-phosphorylated form.

### Crystallization of amino acid 59 mutants of H-RAS and K-RAS

Purification and crystallization of truncated (residues 1-166) and untagged H-RAS^A59T^ and H-RAS^A59E^ was done using a previously published protocol (Johnson et al., 2016). H-RAS^A59T^ bound to GppNHp was concentrated to 12.1mg/mL, flash frozen, and stored at -80°C before crystallization. H-RAS^A59E^ bound to GppNHp and GDP were stored the same way but concentrated to 18.1 and 15.2mg/mL, respectively. Crystals were grown using the hanging drop vapor diffusion method in 24-well plates sealed with Vaseline at 18°C. Mother liquor constitution was unique for each crystal grown, and each crystal was grown against a mother liquor well volume of 402µL. Crystals were harvested at 2-4 weeks and flash frozen using mother liquor with 30% glycerol. The first crystal of H-RAS^A59T^ bound to GppNHp was crystallized in 2µL by 2µL drops using the following reservoir solution: 2.6mM NaCl, 1mM MgCl2, 15.7mM HEPES pH 7.5, 2.5mM DTT, 37.3mM Ca(OAc)2 and 20.5% PEG3350. The second crystal of H-RAS^A59T^ was crystallized in 1µL by 1µL drops using the following reservoir solution: 10mM Mg(OAc)2, 44.8% PEG400, 15.7mM HEPES pH 7.5, 1mM MgCl2, 3.4mM NaCl, and 2.5mM DTT. H-RAS^A59T^ crystals contained one molecule of H-RAS^A59T^ in the asymmetric unit and exhibited P3221 symmetry. H-RAS^A59E^ bound to GDP was crystallized in 1µL by 1µL drops using the following reservoir solution: 2.6mM NaCl, 1mM MgCl2, 15.7mM HEPES pH7.5, 2.5mM DTT, 43.5mM Ca(OAc)2, and 20.5% PEG 3350. H-RAS^A59E^ bound to GppNHp was crystallized in 1µL by 1µL drops using the following reservoir solution: 2.6mM NaCl, 1mM MgCl2, 15.7mM HEPES pH7.5, 2.5mM DTT, 9.95mM Ca(OAc)2, and 19.9% PEG 3350. Both H-RAS^A59E^ crystals contained two molecules in the asymmetric unit, but GppNHp crystals grew with P1211 symmetry while the GDP crystals grew with P1 symmetry. Data collection for all crystals was done on a home source MicroMax007HF with Cu^2+^ anode and tungsten filament, and a R-Axis IV^++^ detector from Rigaku. Indexing, integration and scaling data processing steps were done using the HKL3000 package (Otwinowski and Minor, 1997). For molecular replacement, a previously refined model of wild-type H-RAS (PDB code 1CTQ) was used for both H-RAS^A59T^ and H-RAS^A59E^ crystals bound to GppNHp (Klink and Scheidig, 2010). H-RAS^A59E^ bound to GppNHp was used as starting models for the H-RAS^A59E^ GDP structures. A single round of simulated annealing was performed before refinement. Molecular replacement and structure refinement was done use the PHENIX program (version 1.11.1-2575) (Adams et al., 2010).

Protein for crystallization of K-RAS^A59E^ bound to GDP was acquired by a different purification scheme than above. His-tagged K-RAS4B^A59E^ was overexpressed in *E. coli* BL21 (DE3) and purified in the presence of 20 μM GDP. Briefly, cells were grown at 37°C in TB medium in the presence of 100 μg/ml of ampicillin to an OD of 0.8, cooled to 17°C, induced with 500 μM IPTG, incubated overnight at 17°C, collected by centrifugation, and stored at -80°C. Cell pellets were lysed in buffer A (25 mM HEPES, pH 7.5, 50 mM NaCl, 5 mM MgCl2, 5% glycerol, 20 μM GDP, 7 mM 2-mercaptoethanol, and 20 mM Imidazole), and the resulting lysate was centrifuged at 30,000g for 40 min. Ni-NTA beads (Qiagen) were mixed with cleared lysate for 30 min and washed with buffer A. Beads were transferred to an FPLC-compatible column, and the bound protein was washed further with buffer A for 10 column volumes and eluted with buffer B (25 mM HEPES, pH 7.5, 50 mM NaCl, 5 mM MgCl2, 5% glycerol, 20 μM GDP, 7 mM 2-mercaptoethanol, and 400 mM Imidazole). The eluted sample was concentrated, then 10-fold diluted in buffer C (20 mM Tris, pH 8.0, 50 mM NaCl, 5 mM MgCl2, 5% glycerol, 20 μM GDP, and 1 mM DTT), applied to Mono-Q column (GE healthcare), and eluted by using 50-500mM NaCl gradient. KRAS containing fractions were concentrated and passed through a Superdex 75 10/300 column (GE healthcare) in a buffer containing 25 mM HEPES, pH 7.5, 200 mM NaCl, 5 mM MgCl2, 5% glycerol, 20 μM GDP, and 1 mM DTT. Fractions were pooled, concentrated to approximately 40 mg/ml, and frozen at -80°C. For crystallization, a Formulatrix NT8, RockImager and ArtRobbins Phoenix liquid handler was used to dispense a 100 nl sample of 800 µM K-RAS^A59E^ with an equal volume of 1.5M NaMalonate and 0.1 M Hepes pH 7.5 for crystallization by sitting-drop vapor diffusion. Crystals were grown at 27°C for three days before harvest. Diffraction data were collected at beamline 24ID-E of the NE-CAT at the Advanced Photon Source (Argonne National Laboratory). Data sets were integrated and scaled using XDS (Kabsch, 2010). Structures were solved by molecular replacement using the program Phaser and the search model PDB entry 5TAR (Dharmaiah et al., 2016; McCoy et al., 2007). Iterative manual model building and refinement using Phenix and Coot led to a model with excellent statistics (Adams et al., 2010; Emsley and Cowtan, 2004). Statistics for all crystal data can be found in Table 1.

#### NMR spectroscopy

Isotopically ^15^N labeled K-RAS (residues 1-185 of K-RAS4B with a C118S substitution bearing A59T or A59E substitutions or wild-type A59) was expressed as with an N-terminal His-tag from the pET28 vector in the E. coli strain BL21 (DE3 Codon^+^). A codon-optimized sequence encoding K-RAS4B was synthesized (GenScript) and the A59 codon was mutated using QuikChange site-directed mutagenesis (Agilent). Transformed bacteria was cultured at 37°C in M9 minimal media in the presence of kanamycin and chloramphenicol and supplemented with 1 g/L ^15^N ammonium chloride until the O.D.600 nm reached 0.6. Protein expression was then induced with 0.2 mM IPTG (Isopropyl β-D-1-thiogalactopyranoside) at 15°C overnight.

After centrifugation of the culture, the cell pellets were re-suspended with lysis buffer (5 0mM Tris, 150 mM NaCl, 0.1% NP-40, 10% Glycerol, 10 mM Imidazole, 5 mM MgCl2, 1 mM phenylmethylsulfonyl fluoride (PMSF), 10mM β-mercaptoethanol and lysozyme at pH 8.0) and lysed by sonication. Following centrifugation, His-tagged K-RAS proteins were purified by Ni^2+^-NTA column affinity chromatography from the soluble fraction, the buffer was exchanged to reduce the imidazole concentration, then K-RAS was further purified by size exclusion chromatography (Superdex^TM^ S75 26-60 column, Cytiva) in a running buffer comprising 20 mM HEPES, 100 mM NaCl, 5 mM MgCl2, and 2 mM tris(2-carboxyethyl)phosphine (TCEP), pH 7.4. To prepare GMPPNP-loaded K-RAS, the protein was incubated in the presence of the nucleotide analog, EDTA and calf intestinal phosphatase, which were then removed by passage through a size exclusion desalting column (PD-10, Cytiva).

^1^H-^15^N HSQC spectra were collected with 8 scans at 25°C on a Bruker NEO III HD 800MHz spectrometer equipped with a 5-mm TXO CryoProbe. K-RAS samples were concentrated (wild-type: 500 μM; K-RAS^A59T^: 360 µM; K-RAS^A59E^: 500 µM) in size exclusion buffer plus 5% (vol/vol) D2O. NMR data were processed and analyzed using NMRPipe (Delaglio et al., 1995) and NMRView (Johnson, 2004). Chemical shift perturbations (CSPs, chemical shift changes of the NH cross-peaks of the A59T/E mutants relative to wild-type), were calculated using the formula ΔδNH.N(ppm)=√([(ΔH)]^2^+[(ΔN/5)]^2^) and plotted against residue number using GraphPad 9.1.0. The NH resonances of the mutants were assigned based on previous assignment of wild-type K-RAS (BMRB entry 27720) together with a 3D ^15^N-edited NOESY HSQC spectrum (mixing time 120ms). Chemical shift changes were mapped onto the structure of wild-type K-RAS bound to GDP (PDB 6MBU) as well as our new structure of K-RAS^A59E^ using Chimera 1.15.

### Autophosphorylation and dephosphorylation experiments

Phosphorylated β-casein was used as a positive control for Lambda phosphatase (LMP) activity for dephosphorylation experiments. 90µL of tagged K-RAS^A59T^ at 10mg/mL and 10µL of GTP or GDP at 50mg/mL were incubated overnight at 37°C. After overnight incubation, 10µL of GTP and K-RAS^A59T^ mixture, or β-casein, was incubated at 30°C or 95°C for 10 minutes, and then dephosphorylation was tested by adding 1µL of K-RAS or β-casein to 49µL of 1x phosphatase buffer supplemented with MnCl2 and either 400 units of LMP (New England BioLabs) or water. Dephosphorylation was allowed to proceed for 2, 4, and 6 hours at 30°C. The kinetics of autophosphorylation were also done using purified tagged K-RAS^A59T^ with tagged SOS1 or GAP334. Autophosphorylation reactions were done using a 20X reaction buffer of 1M Tris-HCL and 20mM MgCl2 at pH 7.5. Purified proteins were diluted to initial starting concentrations of 250µM K-RAS^A59T^ and 100µM GAP and SOS1. For the K-RAS^A59T^ reaction without regulators, reactions were run at the following conditions: 2.5µL of reaction buffer, 5µL GTP (25mg/mL) dissolved in water, 1-16µL of 250µM KRAS4B^A59T^ and finally volume balanced to 50µL with water. For autophosphorylation in the presence of GAP and SOS1, the above reaction mixture was used, except 4µL of 250µM K-RAS^A59T^ was fixed for each reaction and 0.5-20µL of purified regulator was added to each reaction and balanced with water to 50µL. Each autophosphorylation experiment was done over 8 hours at 37°C. Each hour, a 2µL sample was taken from each reaction and then added to 2µL of 6X SDS-PAGE sample buffer and 8µL of water. Samples were then boiled for 5 minutes at 70°C. Changes in phosphorylation were detected by 12.5% SDS-PAGE. While the dephosphorylation experiments were done using Coomassie blue staining, the kinetics of autophosphorylation were measured using western blot. Quantification was performed by measuring changes in the upper band versus the sum of the upper and lower bands.

### RAS hydrolysis and nucleotide exchange assays

Assays were done with a BioTeck Synergy microplate reader using black-walled 96-well plates. For GTP hydrolysis, both preloading of GTP and measurement of inorganic phosphate were done in a 96-well plate format. First, a GTP preloading mixture consisting of 72µM K-RAS, 10mM GTP, 4mM EDTA, and 2mM DTT in a final volume of 25µL was stored on ice until use. Second, a GTP hydrolysis mixture consisting of 60mM Tris, 60mM NaCl, 6mM MgCl2, 1.6mM DTT, 240µM of 2-amino-6-mercapto-7-methylpurine riboside (MESG), 1.2 U/mL of purine nucleoside phosphorylase (PNP), and varying concentrations of GAP was brought to a final volume of 125µL. The GTP exchange and reaction mixtures were incubated at 37°C for 35 minutes in the microplate reader in separate wells in a black-walled 96-well plate. After 35 minutes, the GTP preloading mixture was added to the hydrolysis mixture and nucleotide hydrolysis was measured over 90 minutes at 37°C. A change in inorganic phosphate due to GTP hydrolysis was measured by the difference in absorbance at 360nm from a reference reaction without K-RAS protein.

Nucleotide exchange assays were performed for 2’/3’-O-(N-Methyl-anthraniloyl)-guanosine-5’-triphosphate (mant-GTP) or mant-GDP. Mant-GTP and GDP stocks were purchased as 5mM stocks from Invitrogen. In order to measure nucleotide exchange at 37°C, we generated a protein mixture containing 1.6µM K-RAS, 52.5mM Tris pH 7.5, 52.5mM NaCl, 5.25mM MgCl2, 4.2mM DTT and varying concentrations of SOS1 at a final volume of 140µL, and a second mixture containing 225µM of mant-GTP or GDP, 48.6mM Tris, 48.6mM NaCl, 4.9mM MgCl2, 1.9mM DTT at pH7.5 at a final volume of 10µL. The two mixtures were placed in the 96-well plate, wrapped in foil, and incubated in the dark for 30-40 minutes. Once combined, measurement of nucleotide exchange were taken every 65 seconds for 2 hours at 37°C. Measurement of mant-nucleotide association with K-RAS was done by excitation at 360nm, detection of fluorescent emission at 440nm, and subtraction of a reference reaction lacking K-RAS protein. Replicate measurements of hydrolysis and nucleotide exchange were used to determine an apparent first-order rate constant (*k*obs) using the program GraphPad Prism (Notredame et al., 2000).

#### Measurement of K-RAS^A59T^ in cell proliferation experiments

LIM1215, HEK293t, and NIH3T3 cell cultures were obtained from the American Type Culture Collection (ATCC). SNU-175 cells were obtained from the Korean Cell Line Bank (KCLB). SNU-175, LIM1215, HEK293t, and NIH3T3 cells were maintained in accordance ATCC and KCLB protocols, and short tandem repeat (STR) genotyping, performed by LabCorp, was done to ensure cell line authenticity. Measurement of autophosphorylation in response to EGF or insulin signaling was done using SNU-175 cells seeded at a starting density of 7.5x10^5^ cells/well in a 6-well plate, allowed to adhere overnight, and then serum starved for 24 hours. After serum starvation, induction was done by either addition of vehicle (BSA suspended in Hepes), 1.5µg/mL of hEGF, or 5µg/mL of insulin. To measure changes in K-RAS autophosphorylation, cells were lysed at different times. Cell lysate was collected in the following manner. First, cells were gently washed once with 1mL of a phosphate buffered saline solution (PBS). Next, 150µL of radioimmunoprecipitation assay (RIPA) buffer (Boston Bioproducts) was applied to the well and lysis was allowed to proceed for 15 minutes at 4°C with gentle rocking. Cell lysate was collected and clarified by high-speed centrifugation at 4°C for 15 minutes before flash freezing in liquid nitrogen and storage at -80°C. For K-RAS knockdown experiments, shRNA in the pLKO-tet plasmid vector targeting *KRAS*, or GFP as a negative control, was obtained from Addgene (#116871)(Shao et al., 2014). Lentivirus was produced as in Ref. (Shao et al., 2014) with HEK293t cells, and SNU-175 cells were infected for 24 hours and allowed to expand for 1 week before selection with puromycin. For K-RAS knockdown experiments, puromycin selected cells were seeded at a starting density of 5x10^4^ cells/well in a 24-well plate. Cells were allowed to adhere overnight before induction of shRNA against *KRAS* using RPMI 1640 media supplemented with 2µg/mL of doxycycline. To monitor changes in K-RAS autophosphorylation in response to *KRAS* knockdown, cells were lysed every 24 hours using the same method as above. To measure the effect of K-RAS knockdown on SNU-175 cell proliferation, parental SNU-175 cells, or cells infected with shRNA suppressing K-RAS or GFP, were grown for 1 week at 2µg/mL of doxycycline. After one week, 50,000 cells/well were seeded in 24-well plates and allowed to grow for 13 days. A doxycycline positive and negative set of proliferation experiments were performed in parallel.

Ectopic expression of mutant HA-K-RAS4B was done using the pBABE-hygro vector (Addgene plasmid #1765) (Morgenstern and Land, 1990). Production of retrovirus for ectopic expression in NIH3T3 cells was done using the pCL packaging system in HEK293t cells (Naviaux et al., 1996). First, HEK293t cells were lipofectamine transfected at 70% confluence. Virus was collected on the third day and used to infect NIH3T3 cells at 20% confluence overnight. After 48 hours, infected cells were selected using 300µg/mL of hygromycin for one week. Cell lysate for effector binding assays were collected in the following manner. Infected NIH3T3 cells were grown to high density in 175mm dishes, and then washed twice with ice-cold 10mL of PBS and lysed with 400-600µL of MLB solution (25mM HEPES pH 7.5, 150mM NaCl, 1% IGEPAL-CA630, 0.25% sodium deoxycholate, 10% glycerol, and 10mM MgCl2). Lysate and cell debris were immediately scrapped into Eppendorf tubes, rotated for 30 minutes at and then clarified for 30 minutes using high-speed clarification at 4°C. Cell proliferation was measured over four days at the indicated times at 6% FBS and in quadruplicate by seeding 6-well TC culture plates with 5000 cells/well. Cells were collected and counted using a Nexcelom Bioscience Cellometer^®^ AutoT4 cell counter.

#### KRAS-effector binding and precipitation assays

The RAS binding domain of C-RAF kinase (RAF-RBD) (Brtva et al., 1995) and full-length human RASSF5 from the RAS clone collection were obtained from Addgene (Raf1-RBD: #13338, RASSF5: #70545). The RAS clone collection was a gift from Dominic Esposito (Addgene kit 1000000070). GST tagged RAF1-RBD (GST-RAF-RBD) was expressed and purified from BL21 cells as in ref. (Taylor et al., 2001). GST-tagged RASSF5 (GST-RASSF5) sequence was transferred into the pDEST15 expression vector (Thermofisher), and then expressed and purified from BL21-A1cells. Our protocol for measuring effector-K-RAS interactions was done as in REF (Taylor et al., 2001), with the following changes. After protein induction, by either isopropyl β-d-1-thiogalactopyranoside (for GST-RAF-RBD) or arabinose (for GST-RASSF5), cells were washed once in HBS buffer (25mM HEPES pH 7.5, 150mM NaCl), resuspended in RLB solution (20mM HEPES in pH 7.5, 120mM NaCl, 10% glycerol) and then lysed by sonication. Lysate was clarified by high-speed centrifugation at 4°C, flash frozen and stored at -80°C until use. Before measuring mutant HA-K-RAS4B precipitation by RAF-RBD or RASSF5, HA-K-RAS4B in NIH3T3 lysates were preloaded with nucleotide by diluting NIH3T3 lysate in MLB to 2µg/µL in the presence of 1mg/mL of nucleotide, 15mM EDTA, and incubated at 32°C for 30 minutes. After nucleotide preloading, 200µL of nucleotide exchanged lysate was added to an equal volume of MLB buffer containing 10µL of GST-RAF1-RBD or GST-RASSF5 loaded glutathione beads. Binding of K-RAS mutants to the different effectors was allowed to proceed for 2 hours at 4°C with gentle rotation. After the precipitation reaction was complete, beads were washed three times (>20,000xg at 4°C) with equal volumes of MLB. The activated state of HA-K-RAS4B in NIH3T3 cells (i.e. cellular GTP-bound state) was done as above except cell lysates were prepared fresh, not subjected to chemical loading of nucleotide, and precipitation was done in a volume of 300µL. K-RAS-effector precipitation was detected by western blot and detection of HA-tag. Affinity precipitation of other RASSF proteins was done as above for RASSF5 using plasmids provided by Addgene RAS clone collection (RASSF1: #70535, RASSF2: #70539, RASSF3: #70541, RASSF4: #70543, RASSF6: #70547, RASSF7: #70549, RASSF8: #70551, RASSF9: #70553, RASSF10: #70537).

### BRET assay

293FT cells were seeded into 12-well dishes at a concentration of 1x10^5^ cells/well. 16 hours after plating, pCMV5-Venus-K-Ras4B and pLHCX-CMV-Raf-Rluc8 constructs (Terrell and Morrison, 2019) were transfected into cells using a calcium phosphate protocol. A 12-point saturation curve was generated in which the concentration of the energy donor construct (Rluc8) was held constant (62.5 ng) as the concentration of the energy acceptor plasmid (Venus) increased (0-1.0 μg). Cells were collected 48 hours after transfection, washed, and plated in PBS. The Rluc8 cofactor coelenterazine-h was added to a final concentration of 3.375 μM, and the BRET signal read 2 min after addition. The BRET signal was measured at 535 nm (bandwidth 30 nm) on the PHERAstar *Plus* plate reader (BMG Labtech) and the Rluc8 signal was simultaneously measured at 475 nm (bandwidth 30 nm). Venus fluorescence was measured independently using an excitation wavelength of 485 nm (5 nm bandwidth), and the emission spectra measured at 530 nm (5 nm bandwidth) on the Tecan Infinite M1000 plate reader. The BRET value for each data point was calculated by dividing the BRET ratio (BRET/Rluc8) by the background signal. The acceptor/donor ratio was equalized against a control where equal quantities of the Venus and Rluc8 constructs were transfected. Data was analyzed using GraphPad Prism. Non-linear regression was used to plot the best fit hyperbolic curve, and values for BRETmax and BRET50 were obtained from the calculated best fit curves.

### Co-immunoprecipitation experiments

HeLa cells were plated at a concentration of 8 x 10^5^ per 10 cm dish 18-24 hours prior to transfection. pCMV5-Venus-K-RAS constructs were then transfected into cells using the XtremeGENE9 transfection reagent per the manufacturer’s instructions, using a 2:1 ratio of XtremeGENE9 to DNA. 48 hours after transfection, serum-starved cells were washed twice with ice cold PBS and lysed in1% NP-40 lysis buffer (20mM Tris [pH 8.0], 137 mM NaCl, 10% glycerol, 1% NP-40 alternative, 0.15 U/mL aprotinin, 1 mM phenylmethylsulfonyl fluoride, 0.5 mM sodium vanadate, 20 μM leupeptin; 500mL/10 cm dish) for 15 min at 4°C on a rocking platform. Lysates were clarified by centrifugation at 14,000 rpm for 10 min at 4°C, after which the protein content was determined by Bradford assays. Lysates containing equivalent amounts of protein were incubated with either anti-GFP/Venus (rat monoclonal) or anti-C-RAF (mouse monoclonal) and protein G sepharose beads for 2 hours at 4°C on a rocking platform. Complexes were washed extensively with 1% NP-40 buffer and then examined by western blot analysis along with aliquots of equalized total cell lysate.

### Biolayer interferometry

The binding affinities for the RBD of B-RAF and RA domain of RASSF5 of the K-RAS-GMPPNP (wild-type and A59T/E) NMR samples were measured using an Octet RED-384 biolayer interferometry biosensor instrument (BLI, FortéBio, Sartorius) running Octet Data Acquisition 9.0.0.37, and analyzed with FortéBio Data analysis software. The RBD/RA of B-RAF (residues 150 -233) and RASSF5 (residues 199 - 367) were subcloned into pGEX-4T2 (GE Healthcare) to produce N-terminal GST fusion proteins, which were expressed and purified as described previously (Smith and Ikura, 2014). GST-tagged B-RAF-RBD was immobilized by incubating anti-GST-conjugated biosensors (FortéBio) with GST-B-RAF-RBD (10 μg/ml) for 10 mins. The coated sensor was dipped into wells containing a range of concentrations of wild-typ K-RAS, K-RAS^A59T^, or K-RAS^A59E^ (54.7 nM to 20 μM KRAS, as indicated on each sensorgram in Figure S6) in 20 mM HEPES, 100 mM NaCl, 5 mM MgCl2, 2 mM TCEP, 0.5% BSA and 0.05% Tween-20) for 30 or 60 s to monitor association, then transferred to buffer alone to monitor dissociation for 30 or 60 seconds. These binding assays were performed at 25°C in 96-well plates with agitation (1000 rpm). Analogous experiments were performed using GST-RASSF5-RA (immobilized at 5 μg/ml) with K-RAS concentrations ranging from 62 nM to 25 μM K-RAS (as indicated on each sensorgram in Figure S6. Sensors with immobilized GST-RBDs were dipped into buffer alone as a control. Binding data were fitted to a 1:1 binding stoichiometry model using both kinetic and steady state analyses. GraphPad 9.1.0 was used to perform t-tests and prepare graphs.

### Western blotting and antibodies

All samples were run on either homemade 12.5% polyacrylamide gels or criterion pre-cast TGX gels from Bio-Rad. All antibodies were used following manufacturer protocols, and quantitation and analysis were done using a Li-COR Biosciences imaging system. The following primary antibodies were used: Pan-RAS (RAS10, Millipore-Sigma, cat. #05-516), K-RAS (polyclonal KRAS antibody, Proteintech, cat. #12063-1-AP), Vinculin (Vinculin (E1E9V) XP, Cell Signaling Technologies, cat. #13901), MEK1 (MEK1 antibody, BD Biosciences, cat. # 610122), pMEK (Phospho-MEK1/2 (Ser217/221) antibody, Cell Signaling Technologies, cat. #9121), Anti-HA-tag (6E2) (Cell Signaling Technologies, cat. #2367), Anti-GST (91G1) (Cell Signaling Technologies, cat. #2625), pC-Raf (s289/296/301) (Cell Signaling Technologies, cat. #9431), C-Raf (BD Pharmagen, cat. #610152), B-Raf (Santa Cruz, cat. #sc-5284), A-Raf (Santa Cruz, cat. #sc-408), GFP/Venus (MBL International, rat monoclonal, cat. # D153-3), GFP/Venus (Roche, mouse monoclonal, cat. #11814460001).

## Acknowledgements

This work was supported by grants from the National Institutes of Health (R01CA178017 and R01CA232372 to K.M.H.) and an award from the Cancer Research UK Grand Challenge and the Mark Foundation to the SPECIFICANCER team. C.W.J. was supported by postdoctoral fellowship 130428-PF-17-066-01-TBG from the American Cancer Society. Work from the Princess Margaret Cancer Centre was supported by the Canadian Cancer Society Research Institute (Grant # 706696 to M.I.) and the Princess Margaret Cancer Foundation. The NMR spectrometer and Octet biosensor were funded by the Canada Foundation for Innovation (CFI). Work done at the NCI was supported by federal funds from the National Cancer Institute, United States under project number ZIA BC 010329. Synchrotron data collection was based upon research conducted at the Advanced Photon Source on the Northeastern Collaborative Access Team beamlines (NIGMS P41 GM103403). The MicroMax007HF used to collect the X-ray data for H-Ras structures at Northeastern University was purchased in part with funds from the NSF MRI-1228897 grant to C.M.

## Author contributions

C.W.J. and K.M.H. conceived and drafted overall paper. C.W.J. was responsible for crystallization studies of H-RAS^A59T^ and H-RAS^A59E^. H.S, E.A.G, J.L., K.S., and S.D. were responsible for crystallization of K-RAS^A59E^. E.M.T. and D.K.M. designed the BRET and co-immunoprecipitation experiments and interpreted the data. E.M.T. performed BRET and co-immunoprecipitation assays. O.P. and J.A.P. were responsible for mass spectrometry experiments validating autophosphorylation. NMR and Biolayer interferometry experiments were done by F.K., T.G., G. M. C. G., C. B. M., and M.I. A.L. helped with creation of Extended Data table 2. C.W.J. performed all other biochemistry and cell biology experiments, as well as molecular dynamics experiments.

## Supplemental Figures and Table

**Figure S1.**
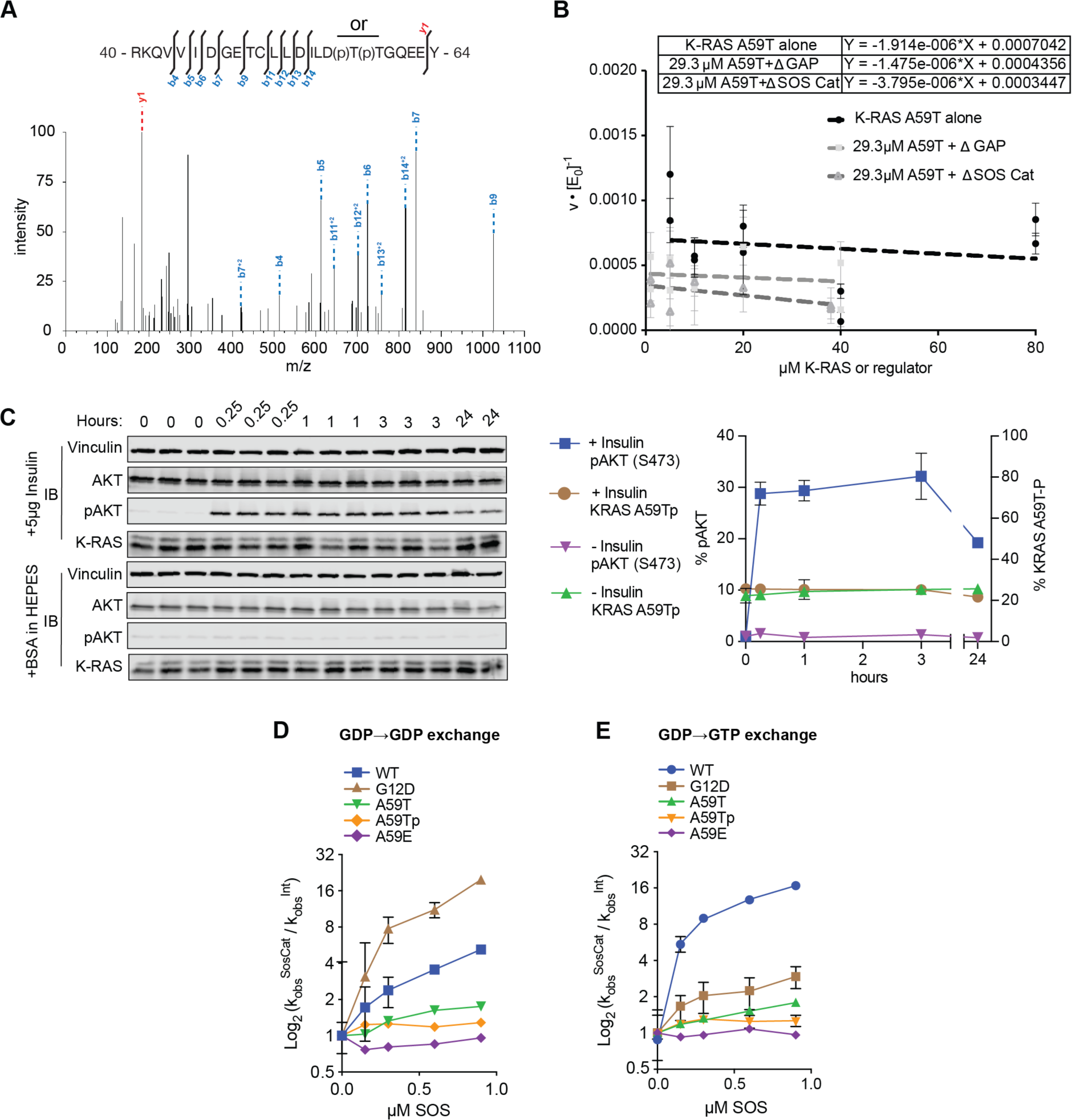
Kinetics of K-RAS autophosphorylation and its effect on nucleotide hydrolysis and exchange. (A) Two precursor peptides, of sequence RKQVVIDGETCLLDILDTTGQEEY, were expected for purified non­phosphorylated (precursor mass: 932.46464, charge: 3+) and phosphorylated (precursor mass: 959.12007, charge 3+) K-RAS^A59T^. Shown is the spectrum for the phosphorylated peptide. (B) Kinetics of Thr59 autophosphorylation in response to increasing concentrations of 6xHis-KRAS4B^A59T^, or increasing concentrations of the catalytic domains of SOS or GAP. (C) SNU-175 cells were serum starved overnight and induced with insulin. Cells were induced at the same time and then stopped at the indicated time points at the top of the blots. Replicates are labelled above the gel and quantification is shown on the right. (D) and (E) Effects of SOS on nucleotide cycling of (E) GDP for mant-GDP and (F) GDP for mant-GTP. Data are depicted as a fold-change relative to *k* of the intrinsic exchange reaction, and data points represent average *k* assuming a first order mechanism (n = 2-4).

**Figure S2.**
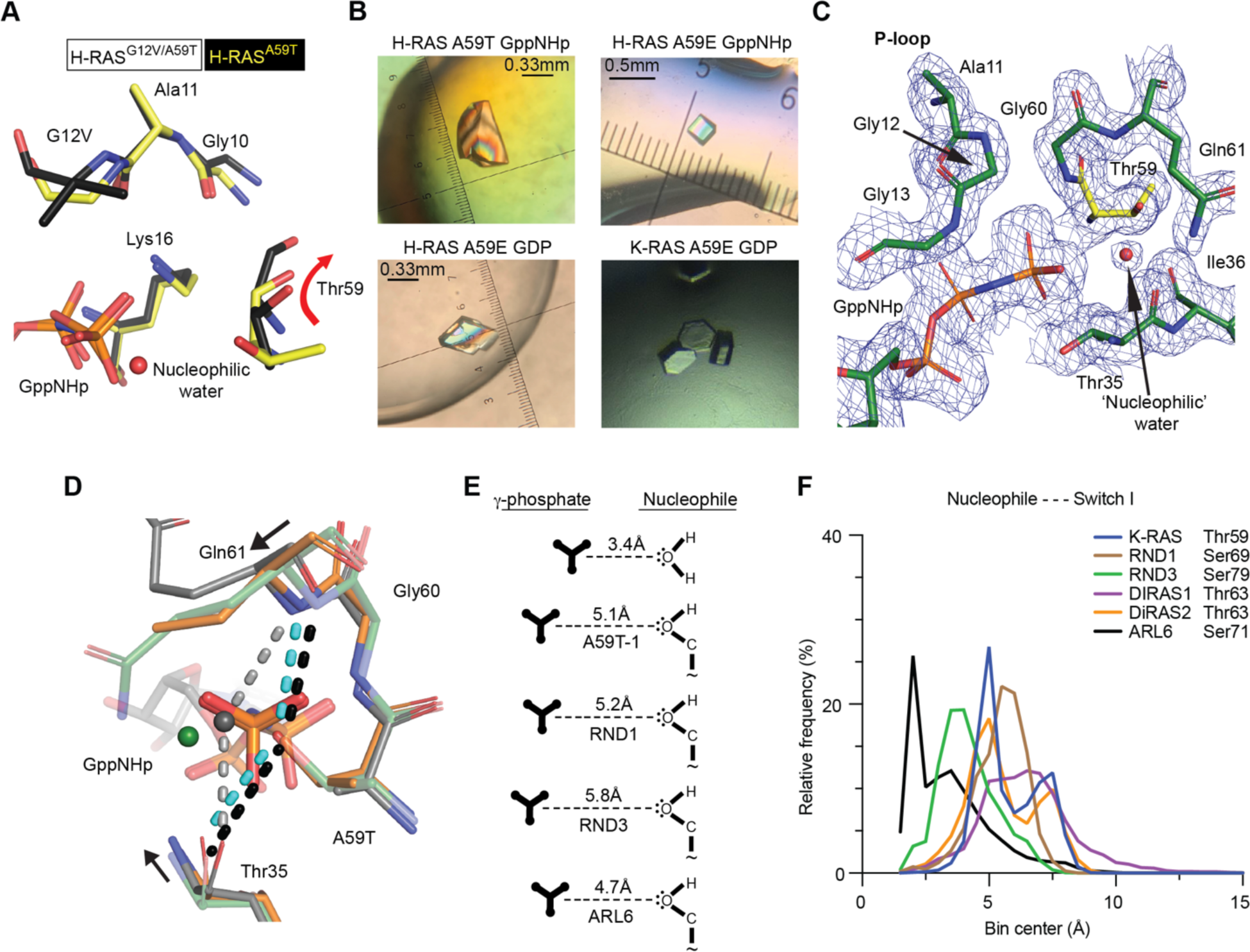
Crystal and dynamic data for autophosphorylation mechanism. (A) Crystal structures of H-RAS^A59T^ (yellow) are different from a previously deposited crystal structure of H-RAs^G12V/A59T^ (black, PDB code 521P) in that T59 is oriented away from the GTP in the G12V/A59T structure. (B) Protein crystals of H-RAS and K-RAS. (C) Active site electron density of H-RAS^A59T^ (sigma = 1.0). (D) Comparison of H-RAS^A59T^ crystal 1 (yellow) and crystal 2 (orange) active sites. The nucleophilic water is colored by crystal. H-RAS^A59T^ crystal 2 shifts toward active site which forces the nucleophilic water molecule to be absent in this crystal. (E) Comparison of T59 orientiation (right) and distance from the g-phosphate of GTP (left) in various GTPases. (F) Nucleophile to switch Iresidue (labeled) distances in small GTPases during MD simualtions.

**Figure S3.**
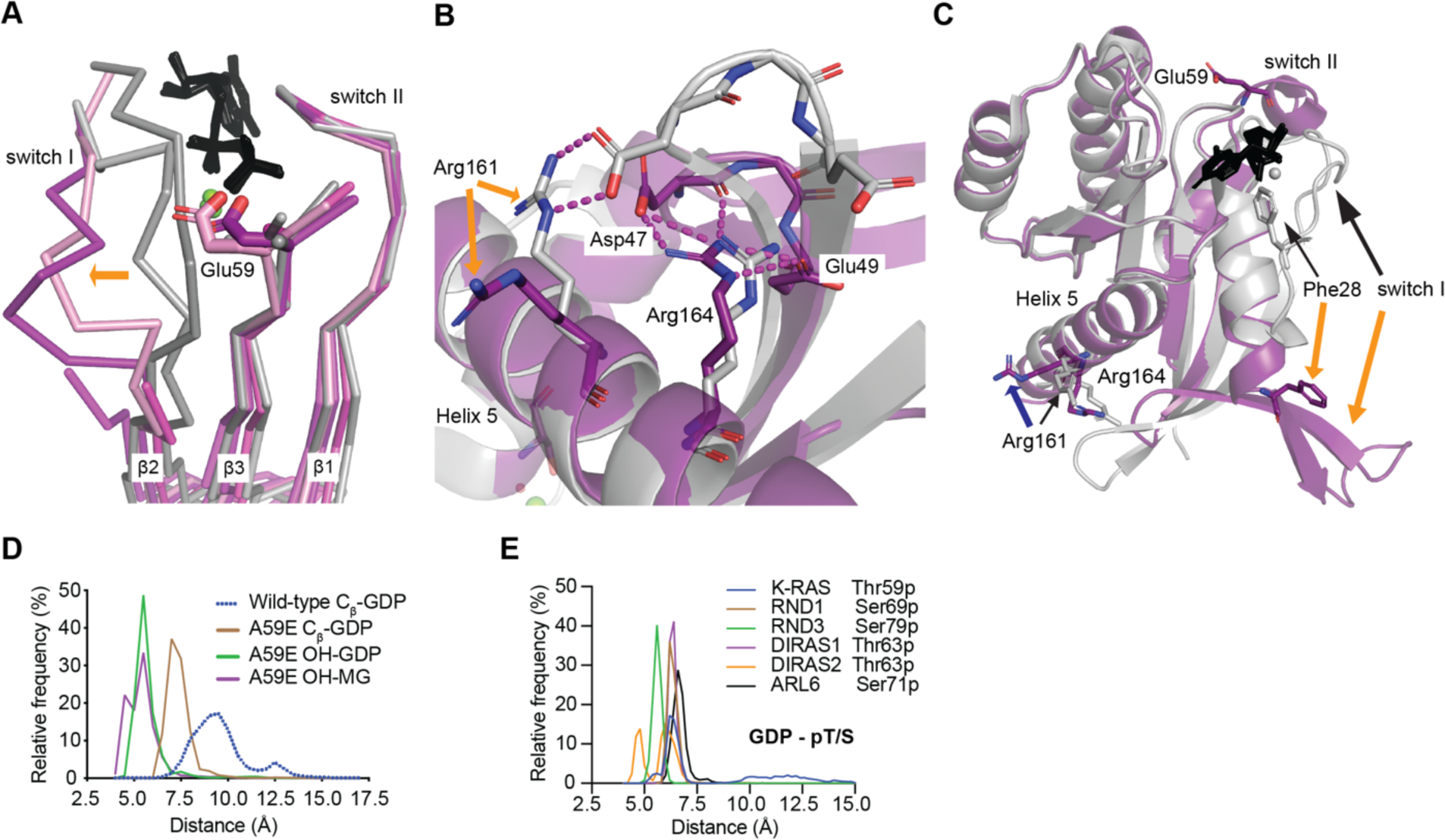
RAS A59E and A59Tp openthe active site andincrease dynamics. (A) H-RAS^A59E^ crystal structures (magenta, purple) show that E59 breaks secondary structure interactions made between the β2 and β3. (B) The switch Iconformation of K-RAS^A59E^ GDP is stabilized by alternative salt-bridge interactions made between R164, D48, E49 and the β-turn connecting β2 and β3. Note that D47 trades its salt-bridge interaction with R161 in WT K-RAS (PDB code 40BE,gray) for R164. (C) The alternate switch I conformation removes the F28-K147 stacking interaction that helps stabilize binding of the guanosine base of GDP. The nucleotide is shown as black sticks on each panel. (D-E) Frequency histograms of bond distances during MD simulations of GDP bound K-RAS mutants and other GTPases phosphorylated by *in silico* modification.

**Figure S4.**
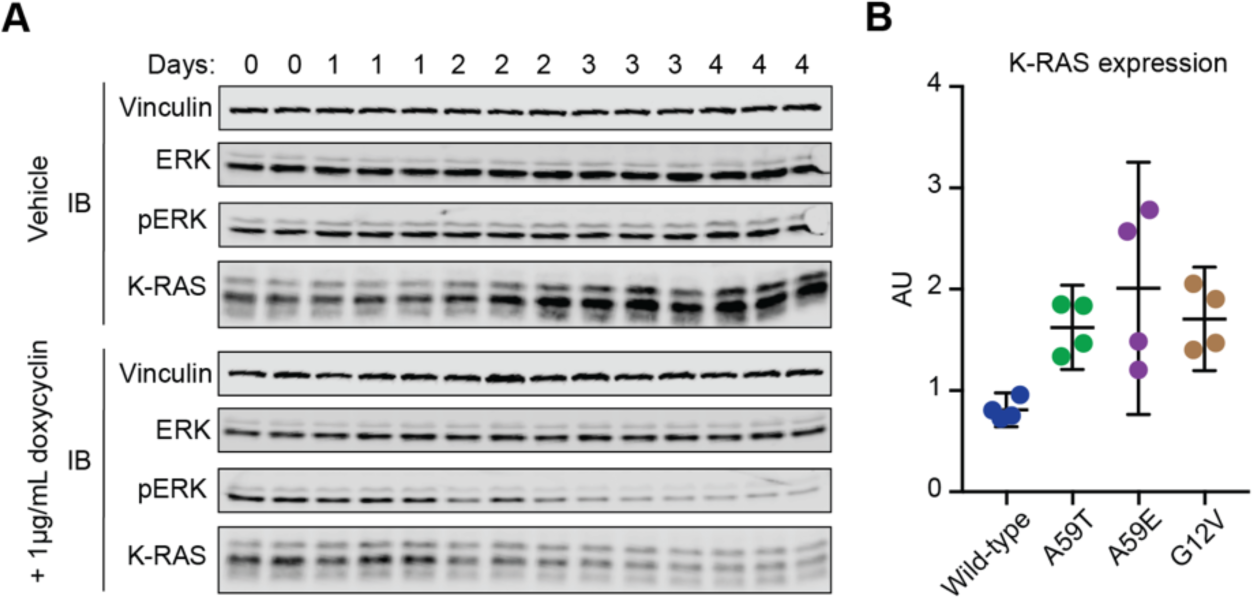
Role of K-RAS^A59T^ and K-RAS^A59E^ in cell transformation. (A) The effect of K-RAS knockdown on the phosphorylated state of K-RAS^A59T^ after induction of shRNA by 1µg/ml of doxycyclin was measured in replicates over the indicated days. (B) Quantification of western blot for HA-tagged K-RAS4B in NIH3T3 cells (n=4). Consistent with our SNU-175 data,a quarter of the ectopically expressed K-RAS^A59T^ was phosphorylated (20.9±4.2%)

**Figure S5.**
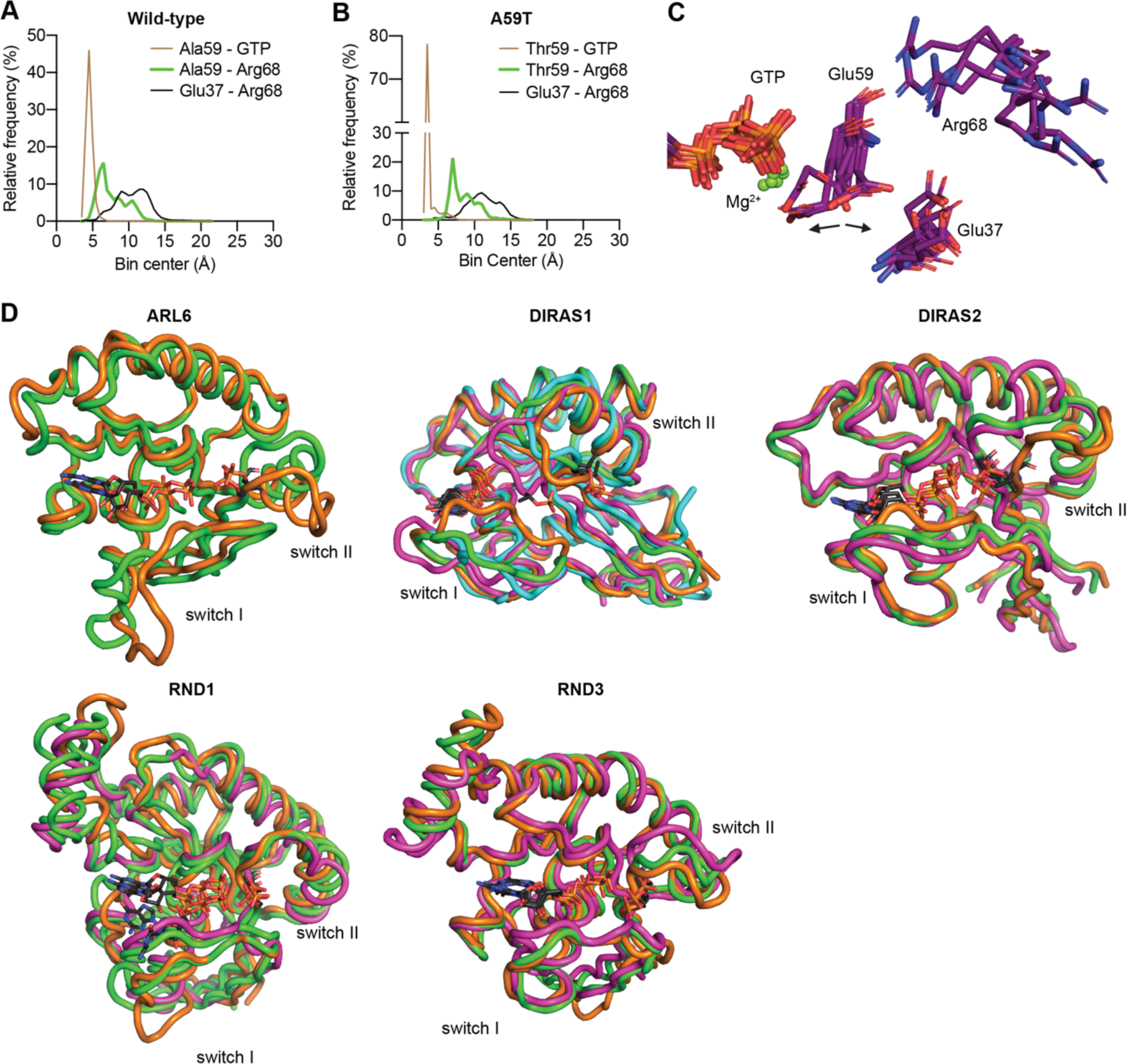
Dynamics of phosphorylated or phospho-mimetic residue 59. (A) and (B), Frequency histograms of bond distances during MD simulations of GDP bound K-RAS mutants and auto­ phosphorylating GTPases phosphorylated b in silico modification. (C) Simulation cluster analysis of K-RAS^A59E^ bound to GTP with a 0.2nm cutoff distance between frames. (D) Cluster analysis of simulations of aGTPase structures starting in the GTP bound state or exchanged to GTP, and with residue 59 modified with phosphorylation in silico.Different colors represent different states from cluster analyses. GTPases are labelled in each panel.

**Figure S6.**
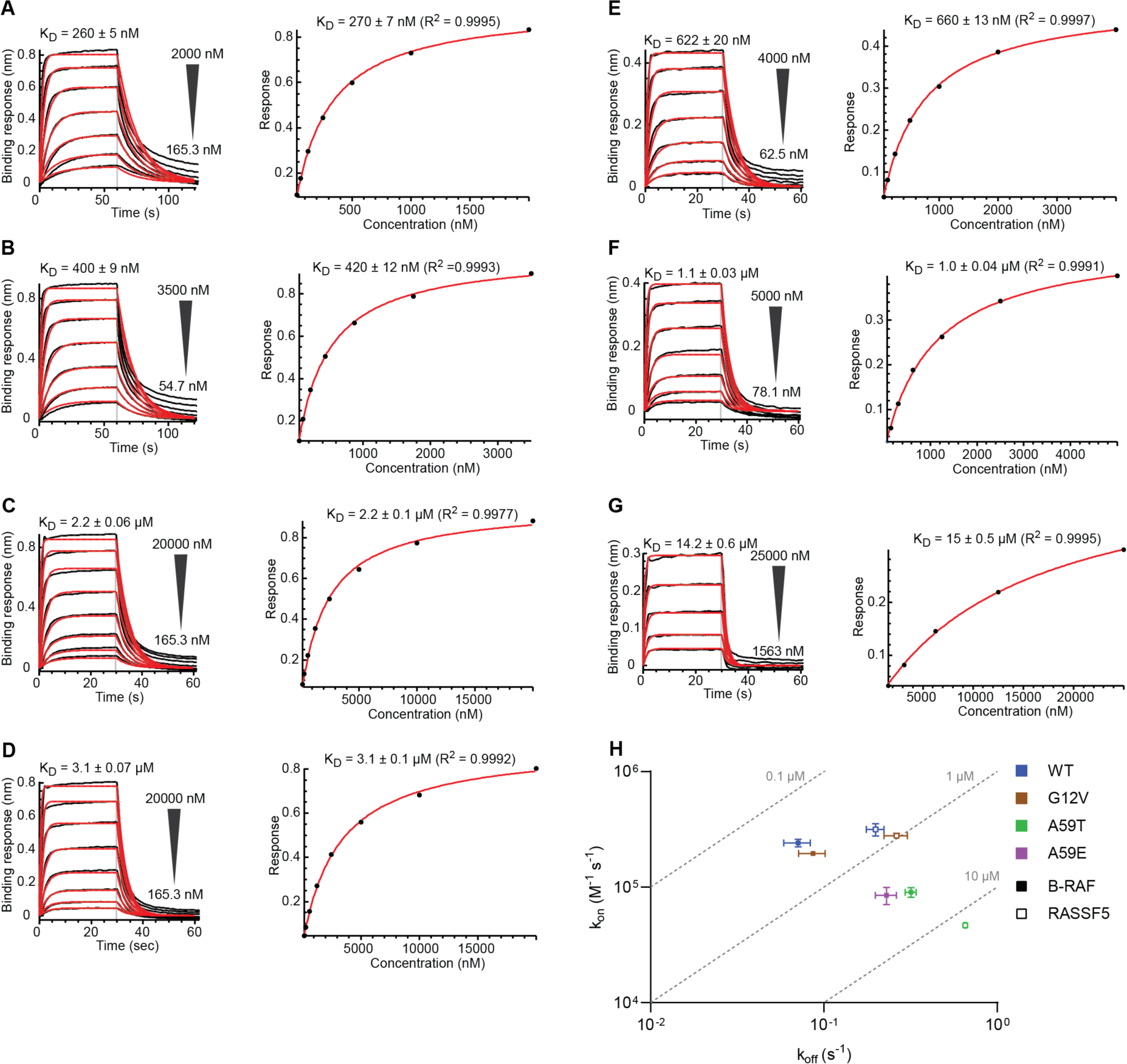
BLI data for B-RAF and RASSF5 interactions with K-RAS proteins. BLI sensorgrams representing interactions between immobilized GST-B-RAF-RBD and GppNHp bound wild-type K-RAS (A), G12V (B), A59T (C), A59E (D), as well as interactions between GST-RASSSF5-RA and GppNHp bound wild-type K-RAS (E), G12V (F) and A59T (G). The interaction between RASSF5 RBD and KRAS A59E was negligible and therefore not shown. The ranges of analyte (KRAS) concentrations are indicated beside each sensorgram. Left panels: sensorgram curves are shown in black with the fitted curves in red and the KD values determined by kinetic analyses displayed above. Right panels: SPR response at equilibrium versus KRAS concentration with KD values determined by steady-state analyses. (H) Iso-affinity plot of the association (k_on_) and dissociation (k_off_) rates derived from BLI kinetic analyses of the interaction between GppNHp bound wild-type K-RAS and mutants G12V, A59T and A59E with B-RAF-RBD (closed symbols) and RASSF5-RBD (open symbols). Kinetic constants were determined by fitting sensograms from a titration series. The grey dashed lines connect points with equal K_D_ values, as indicated. Where no error bars are shown they are smaller than the symbol size (n = 2-4)

**Figure S7.**
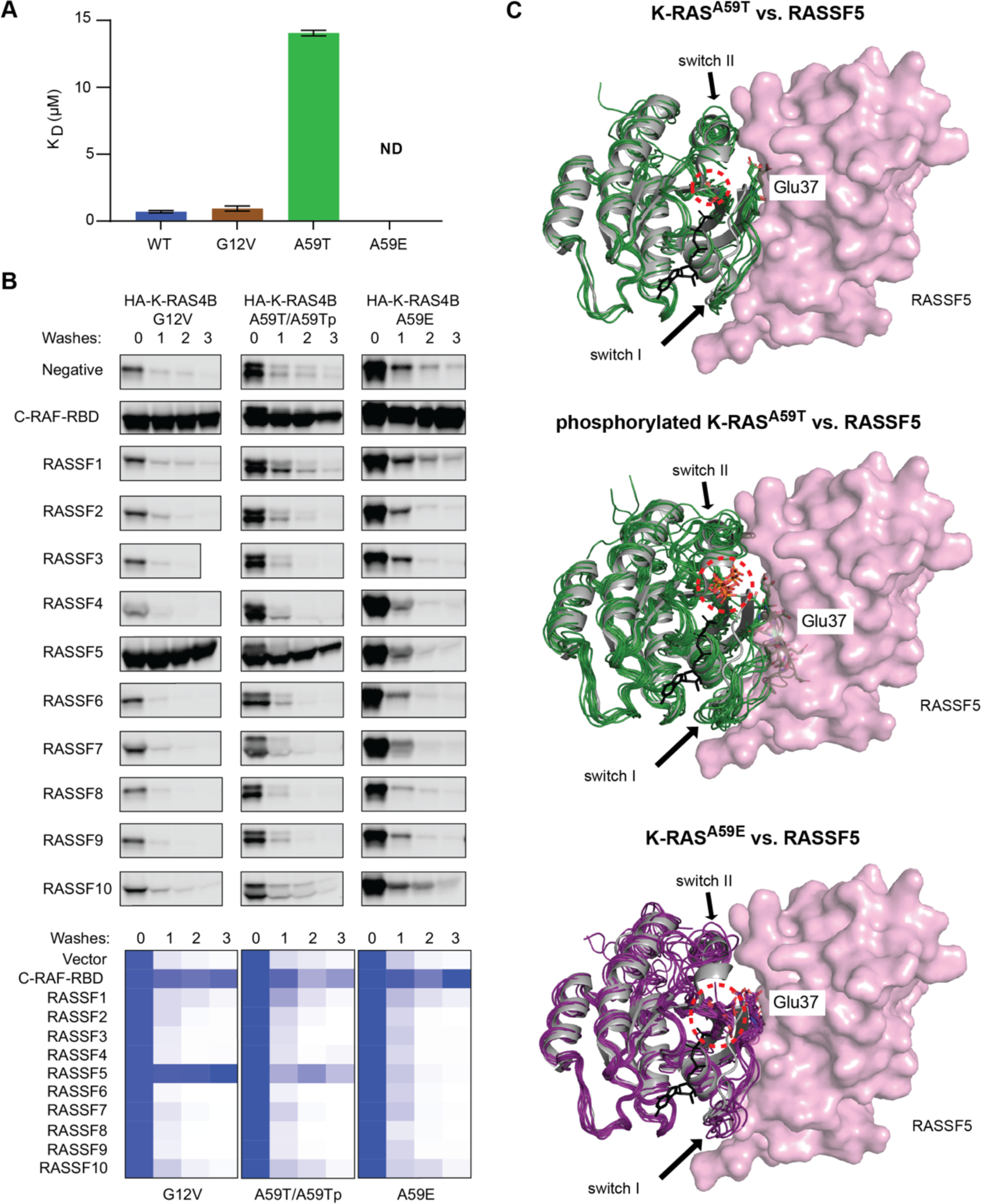
Interaction of phosphorylated K-RAS^A59T^ and K-RAS^A59E^ with RASSF proteins. (A) Affinities of GppNHp bound K-RAS (wild-type,G12V and A59T) for the RA domain of RASSF5 determined by BLI.K-RAS^A59E^ exhibited negligible binding thus a K_D_ value could not be determined. Error bars represent standard deviation (n = 2-4). (B) Precipitation experiments testing the affinity of GTP bound HA-KRAS4B and its mutants to different RASSF proteins. Experiments were done as in Figure S8. Quantification of precipitations were scaled to the ‘0’ wash step (supernatent removal after precipitation, without buffer wash). Note that binding in the K-RAS^A59T^ precipitation set is the combination of both upper and lower bands in the western blot. (C) For each panel, thewild-type crystalstructure of H-RAS is shown as a gray ribbon while simulations are colored strings derived from the same cluster analyses of simulations desribed in Figure 5I. RASSF5-RA domains are shown as colored surfaces. Dotted circles show location of mutation or autophosphorylation.

**Table S1.**
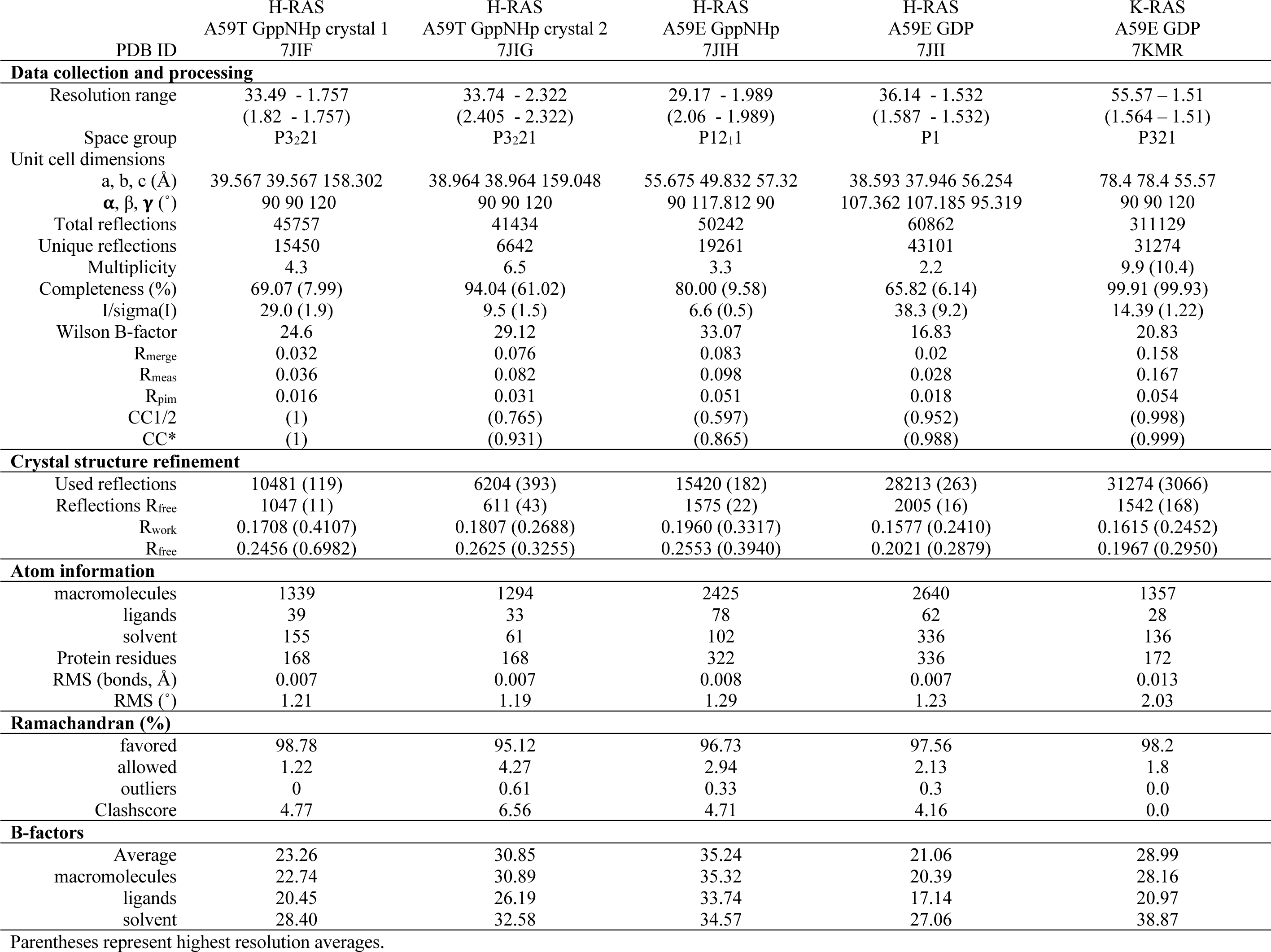
Data collection and structure refinement statistics.

