## Supplemental Figures and Tables for "Regulation of GTPase function by autophosphorylation"

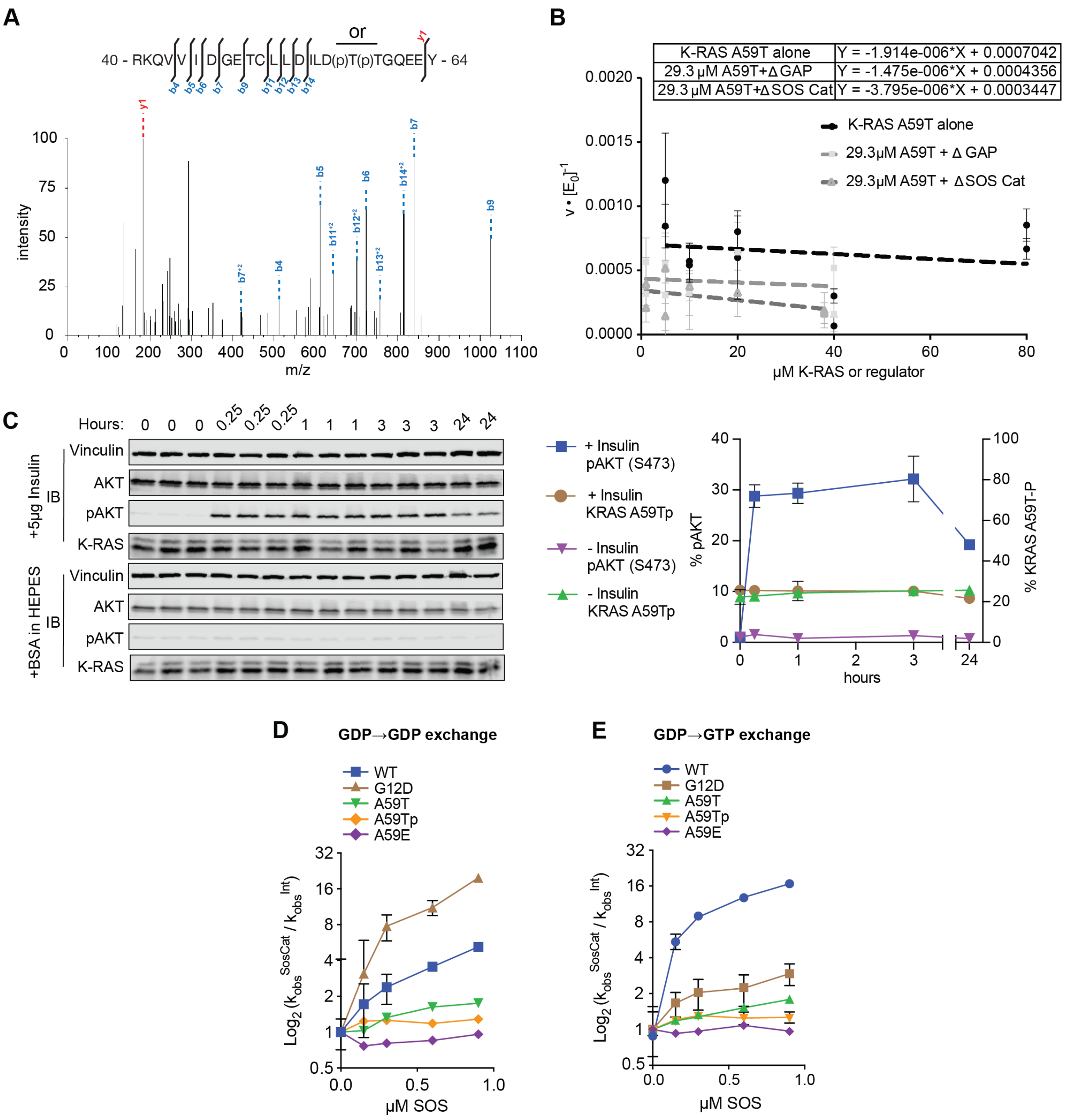


**Figure S1. Kinetics of K-RAS autophosphorylation and its effect on nucleotide hydrolysis and exchange.**

(A) Two precursor peptides, of sequence RKQVVIDGETCLLDILDTTGQEEY, were expected for purified non-phosphorylated (precursor mass: 932.46464, charge: 3+) and phosphorylated (precursor mass: 959.12007, charge 3+) K-RAS$^{A59T}$. Shown is the spectrum for the phosphorylated peptide.

(B) Kinetics of Thr59 autophosphorylation in response to increasing concentrations of 6xHis-KRAS4B$^{A59T}$, or increasing concentrations of the catalytic domains of SOS or GAP.

(C) SNU-175 cells were serum starved overnight and induced with insulin. Cells were induced at the same time and then stopped at the indicated time points at the top of the blots. Replicates are labelled above the gel and quantification is shown on the right.

(D) and (E) Effects of SOS on nucleotide cycling of (E) GDP for mant-GDP and (F) GDP for mant-GTP. Data are depicted as a fold-change relative to *k* of the intrinsic exchange reaction, and data points represent average *k* assuming a first order mechanism (n = 2-4).

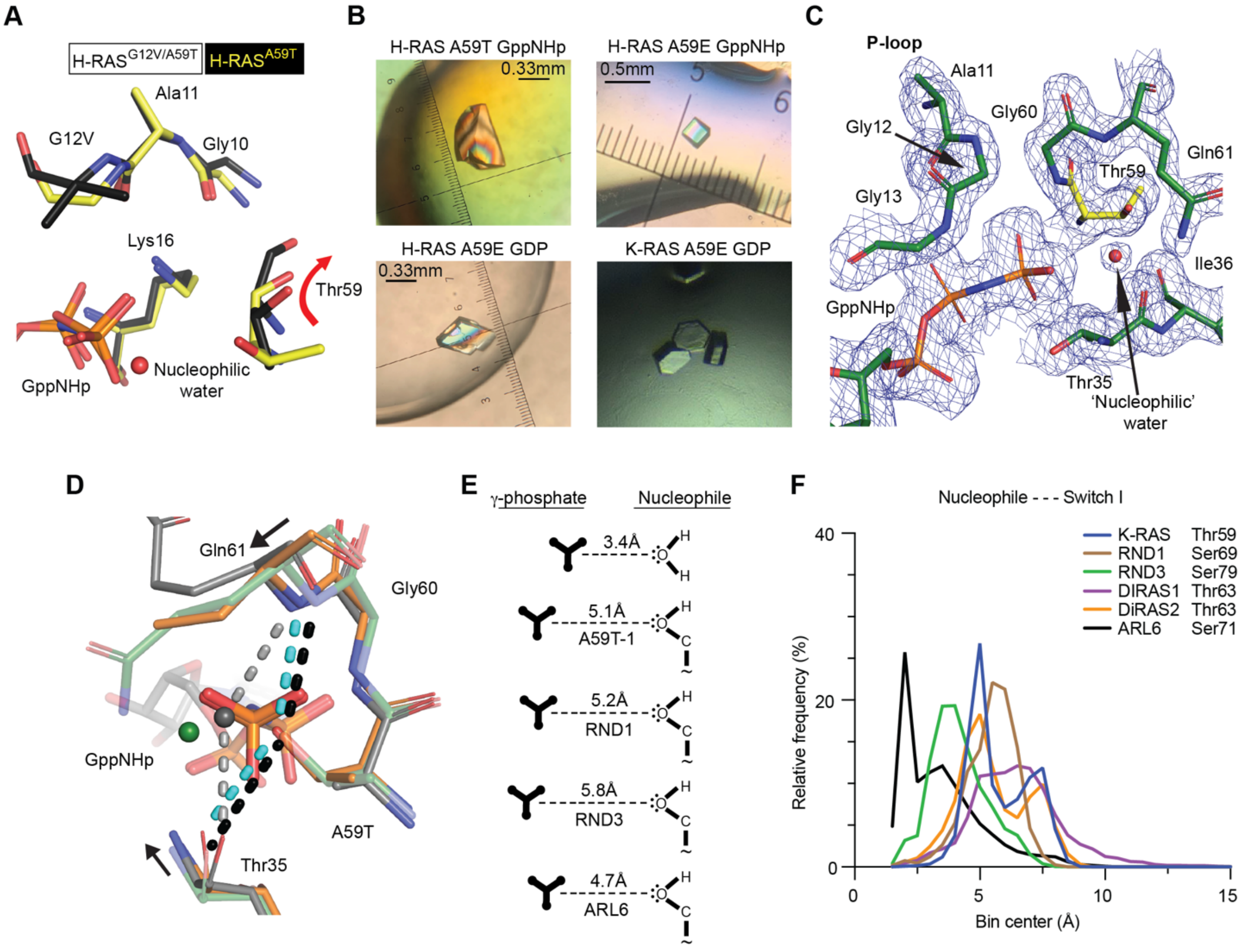


**Figure S2. Crystal and dynamic data for autophosphorylation mechanism.**
(A) Crystal structures of H-RAS$^{A59T}$ (yellow) are different from a previously deposited crystal structure of H-RAS$^{G12V/A59T}$ (black, PDB code 521P) in that T59 is oriented away from the GTP in the G12V/A59T structure.
(B) Protein crystals of H-RAS and K-RAS.
(C) Active site electron density of H-RAS$^{A59T}$ (sigma = 1.0).
(D) Comparison of H-RAS$^{A59T}$ crystal 1 (yellow) and crystal 2 (orange) active sites. The nucleophilic water is colored by crystal. H-RAS$^{A59T}$ crystal 2 shifts toward active site which forces the nucleophilic water molecule to be absent in this crystal.
(E) Comparison of T59 orientiation (right) and distance from the g-phosphate of GTP (left) in various GTPases.
(F) Nucleophile to switch I residue (labeled) distances in small GTPases during MD simualtions.

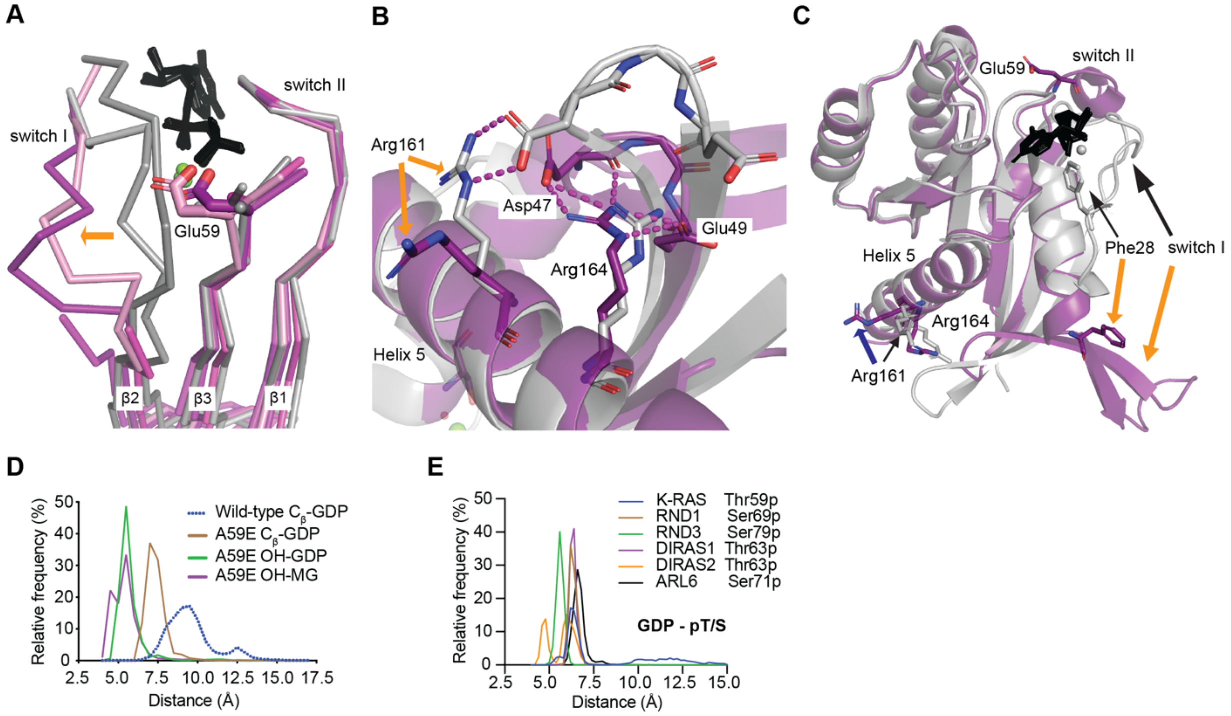


**Figure S3. RAS A59E and A59Tp open the active site and increase dynamics.**
(A) H-RAS$^{A59E}$ crystal structures (magenta, purple) show that E59 breaks secondary structure interactions made between the β2 and β3.
(B) The switch I conformation of K-RAS$^{A59E}$ GDP is stabilized by alternative salt-bridge interactions made between R164, D48, E49 and the β-turn connecting β2 and β3. Note that D47 trades its salt-bridge interaction with R161 in WT K-RAS (PDB code 4OBE, gray) for R164.
(C) The alternate switch I conformation removes the F28-K147 stacking interaction that helps stabilize binding of the guanosine base of GDP. The nucleotide is shown as black sticks on each panel.
(D-E) Frequency histograms of bond distances during MD simulations of GDP bound K-RAS mutants and other GTPases phosphorylated by *in silico* modification.

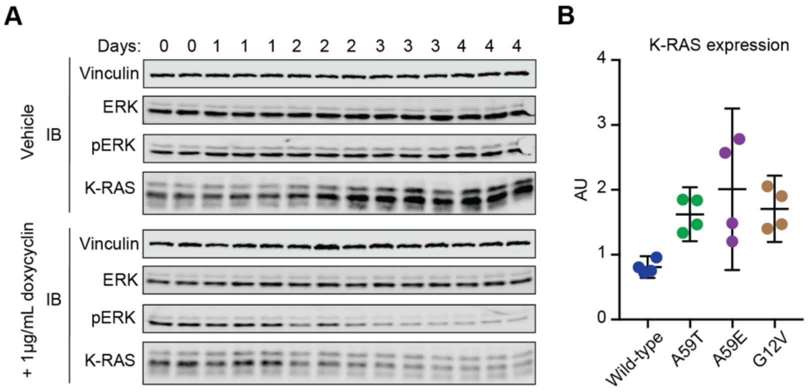


**Figure S4. Role of K-RAS$^{A59T}$ and K-RAS$^{A59E}$ in cell transformation.**
(A) The effect of K-RAS knockdown on the phosphorylated state of K-RAS$^{A59T}$ after induction of shRNA by 1µg/mL of doxycyclin was measured in replicates over the indicated days.
(B) Quantification of western blot for HA-tagged K-RAS4B in NIH3T3 cells (n=4). Consistent with our SNU-175 data, a quarter of the ectopically expressed K-RAS$^{A59T}$ was phosphorylated (20.9±4.2%)

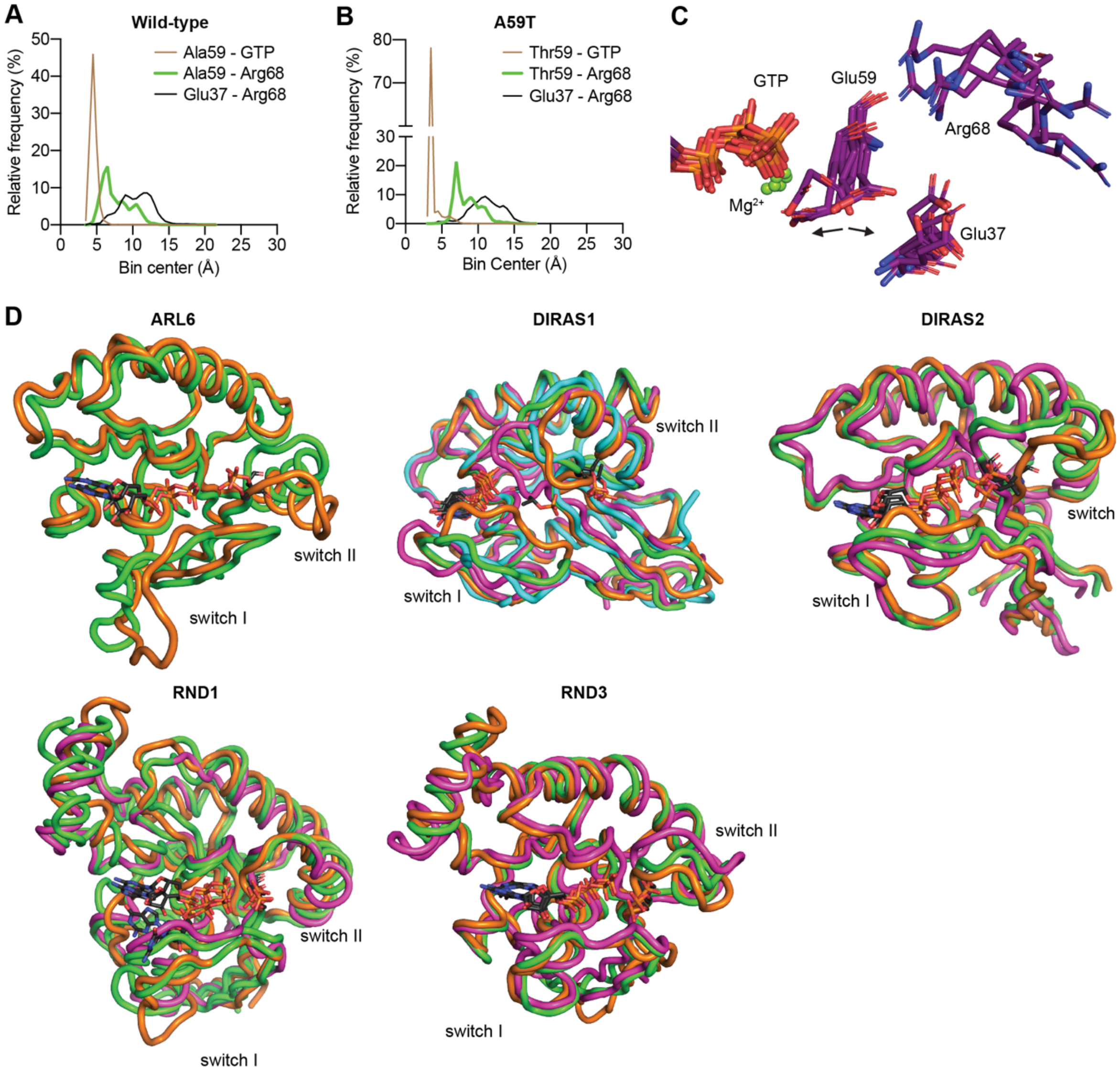


**Figure S5. Dynamics of phosphorylated or phospho-mimetic residue 59.**

(A) and (B), Frequency histograms of bond distances during MD simulations of GDP bound K-RAS mutants and auto-phosphorylating GTPases phosphorylated by in silico modification.

(C) Simulation cluster analysis of K-RAS$^{A59E}$ bound to GTP with a 0.2nm cutoff distance between frames.

(D) Cluster analysis of simulations of aGTPase structures starting in the GTP bound state or exchanged to GTP, and with residue 59 modified with phosphorylation in silico. Different colors represent different states from cluster analyses. GTPases are labelled in each panel.

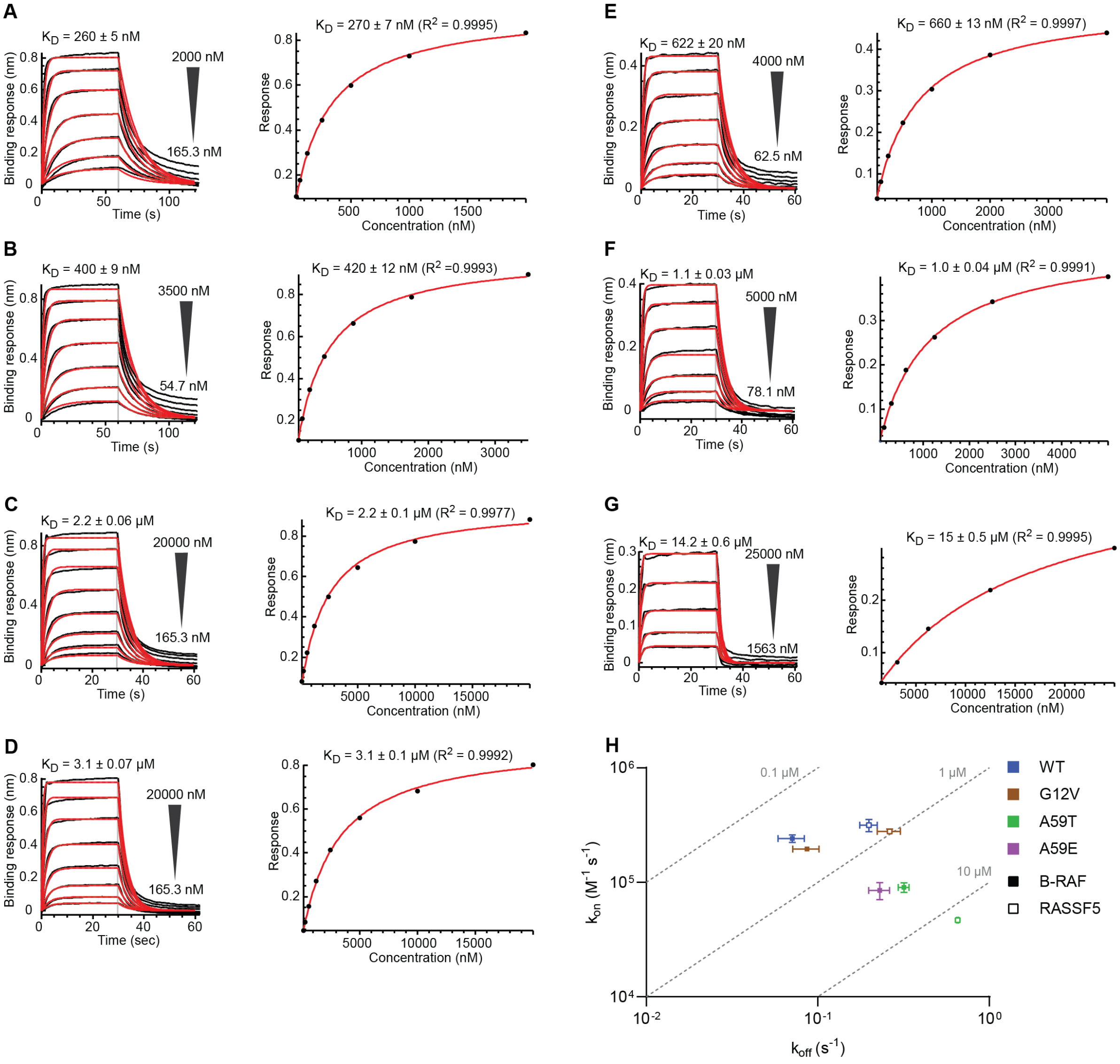


**Figure S6. BLI data for B-RAF and RASSF5 interactions with K-RAS proteins.**
BLI sensorgrams representing interactions between immobilized GST-B-RAF-RBD and GppNHp bound wild-type K-RAS (A), G12V (B), A59T (C), A59E (D), as well as interactions between GST-RASSSF5-RA and GppNHp bound wild-type K-RAS (E), G12V (F) and A59T (G). The interaction between RASSF5 RBD and KRAS A59E was negligible and therefore not shown. The ranges of analyte (KRAS) concentrations are indicated beside each sensorgram. Left panels: sensorgram curves are shown in black with the fitted curves in red and the KD values determined by kinetic analyses displayed above. Right panels: SPR response at equilibrium versus KRAS concentration with KD values determined by steady-state analyses.
(H) Iso-affinity plot of the association ($k_{on}$) and dissociation ($k_{off}$) rates derived from BLI kinetic analyses of the interaction between GppNHp bound wild-type K-RAS and mutants G12V, A59T and A59E with B-RAF-RBD (closed symbols) and RASSF5-RBD (open symbols). Kinetic constants were determined by fitting sensograms from a titration series. The grey dashed lines connect points with equal $K_D$ values, as indicated. Where no error bars are shown they are smaller than the symbol size (n = 2-4)

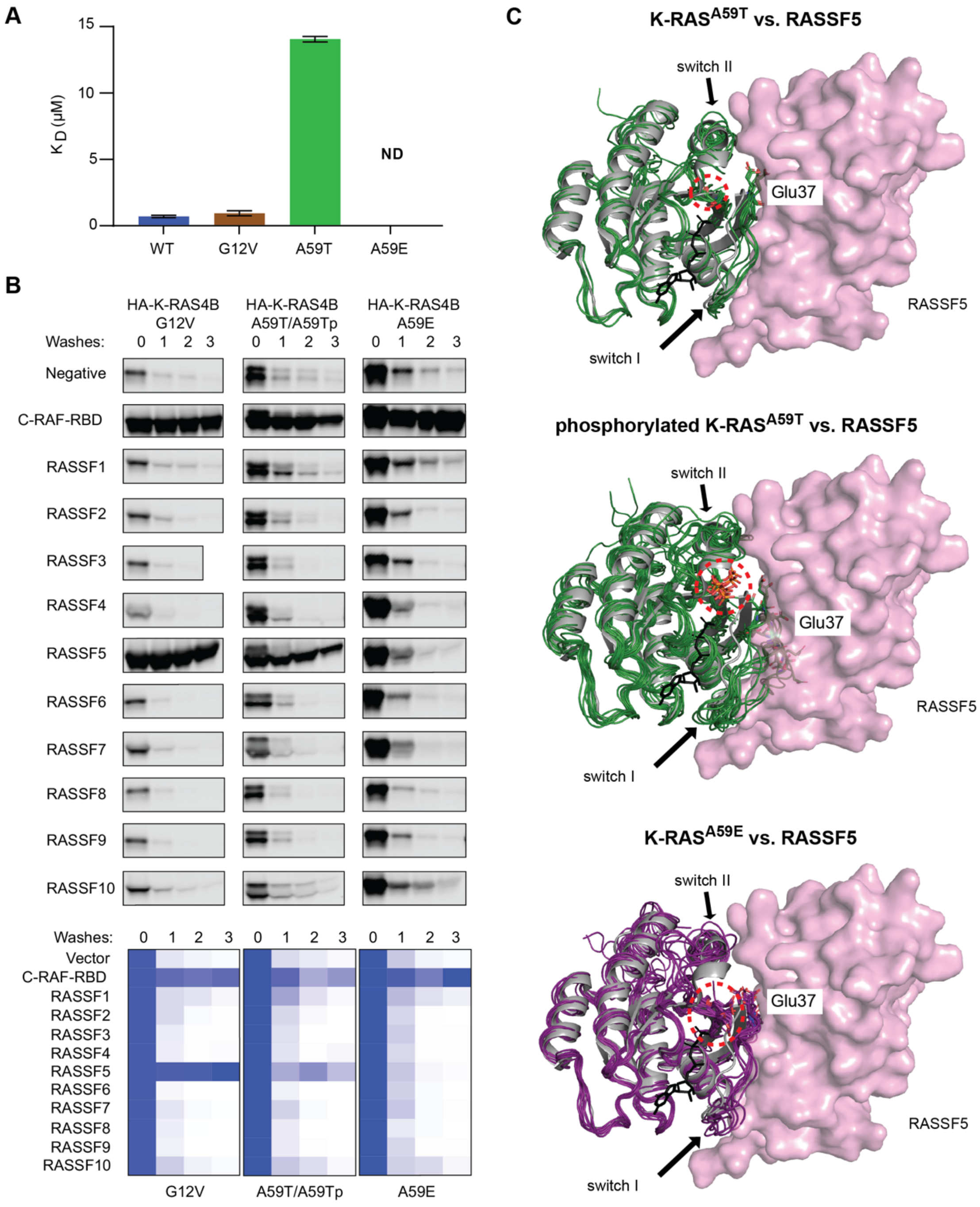


**Figure S7. Interaction of phosphorylated K-RAS$^{A59T}$ and K-RAS$^{A59E}$ with RASSF proteins**.
(A) Affinities of GppNHp bound K-RAS (wild-type, G12V and A59T) for the RA domain of RASSF5 determined by BLI. K-RAS$^{A59E}$ exhibited negligible binding thus a $K_D$ value could not be determined. Error bars represent standard deviation (n = 2-4).
(B) Precipitation experiments testing the affinity of GTP bound HA-KRAS4B and its mutants to different RASSF proteins. Experiments were done as in Figure 5B. Quantification of precipitations were scaled to the '0' wash step (supernatent removal after precipitation, without buffer wash). Note that binding in the K-RAS$^{A59T}$ precipitation set is the combination of both upper and lower bands in the western blot.
(C) For each panel, the wild-type crystal structure of H-RAS is shown as a gray ribbon while simulations are colored strings derived from the same cluster analyses of simulations desribed in Figure 5I. RASSF5-RA domains are shown as colored surfaces. Dotted circles show location of mutation or autophosphorylation.

**Table S1. Data collection and structure refinement statistics.**

| | H-RAS A59T GppNHp crystal 1 | H-RAS A59T GppNHp crystal 2 | H-RAS A59E GppNHp | H-RAS A59E GDP | K-RAS A59E GDP |
|---|---|---|---|---|---|
| PDB ID | 7JIF | 7JIG | 7JIH | 7JII | 7KMR |
| **Data collection and processing** | | | | | |
| Resolution range | 33.49 - 1.757 (1.82 - 1.757) | 33.74 - 2.322 (2.405 - 2.322) | 29.17 - 1.989 (2.06 - 1.989) | 36.14 - 1.532 (1.587 - 1.532) | 55.57 – 1.51 (1.564 – 1.51) |
| Space group | $P3_221$ | $P3_221$ | $P12_11$ | P1 | P321 |
| Unit cell dimensions | | | | | |
| a, b, c (Å) | 39.567 39.567 158.302 | 38.964 38.964 159.048 | 55.675 49.832 57.32 | 38.593 37.946 56.254 | 78.4 78.4 55.57 |
| $\alpha$, $\beta$, $\gamma$ (˚) | 90 90 120 | 90 90 120 | 90 117.812 90 | 107.362 107.185 95.319 | 90 90 120 |
| Total reflections | 45757 | 41434 | 50242 | 60862 | 311129 |
| Unique reflections | 15450 | 6642 | 19261 | 43101 | 31274 |
| Multiplicity | 4.3 | 6.5 | 3.3 | 2.2 | 9.9 (10.4) |
| Completeness (%) | 69.07 (7.99) | 94.04 (61.02) | 80.00 (9.58) | 65.82 (6.14) | 99.91 (99.93) |
| I/sigma(I) | 29.0 (1.9) | 9.5 (1.5) | 6.6 (0.5) | 38.3 (9.2) | 14.39 (1.22) |
| Wilson B-factor | 24.6 | 29.12 | 33.07 | 16.83 | 20.83 |
| $R_{merge}$ | 0.032 | 0.076 | 0.083 | 0.02 | 0.158 |
| $R_{meas}$ | 0.036 | 0.082 | 0.098 | 0.028 | 0.167 |
| $R_{pim}$ | 0.016 | 0.031 | 0.051 | 0.018 | 0.054 |
| CC1/2 | (1) | (0.765) | (0.597) | (0.952) | (0.998) |
| CC* | (1) | (0.931) | (0.865) | (0.988) | (0.999) |
| **Crystal structure refinement** | | | | | |
| Used reflections | 10481 (119) | 6204 (393) | 15420 (182) | 28213 (263) | 31274 (3066) |
| Reflections $R_{free}$ | 1047 (11) | 611 (43) | 1575 (22) | 2005 (16) | 1542 (168) |
| $R_{work}$ | 0.1708 (0.4107) | 0.1807 (0.2688) | 0.1960 (0.3317) | 0.1577 (0.2410) | 0.1615 (0.2452) |
| $R_{free}$ | 0.2456 (0.6982) | 0.2625 (0.3255) | 0.2553 (0.3940) | 0.2021 (0.2879) | 0.1967 (0.2950) |
| **Atom information** | | | | | |
| macromolecules | 1339 | 1294 | 2425 | 2640 | 1357 |
| ligands | 39 | 33 | 78 | 62 | 28 |
| solvent | 155 | 61 | 102 | 336 | 136 |
| Protein residues | 168 | 168 | 322 | 336 | 172 |
| RMS (bonds, Å) | 0.007 | 0.007 | 0.008 | 0.007 | 0.013 |
| RMS (˚) | 1.21 | 1.19 | 1.29 | 1.23 | 2.03 |
| **Ramachandran (%)** | | | | | |
| favored | 98.78 | 95.12 | 96.73 | 97.56 | 98.2 |
| allowed | 1.22 | 4.27 | 2.94 | 2.13 | 1.8 |
| outliers | 0 | 0.61 | 0.33 | 0.3 | 0.0 |
| Clashscore | 4.77 | 6.56 | 4.71 | 4.16 | 0.0 |
| **B-factors** | | | | | |
| Average | 23.26 | 30.85 | 35.24 | 21.06 | 28.99 |
| macromolecules | 22.74 | 30.89 | 35.32 | 20.39 | 28.16 |
| ligands | 20.45 | 26.19 | 33.74 | 17.14 | 20.97 |
| solvent | 28.40 | 32.58 | 34.57 | 27.06 | 38.87 |

Parentheses represent highest resolution averages.
